# LncRNAs interacting with the translation machinery contribute to human neuronal differentiation

**DOI:** 10.1101/2020.10.01.321919

**Authors:** Katerina Douka, Isabel Birds, Dapeng Wang, Sophie Clayton, Abigail Byford, Elton J. R. Vasconcelos, Mary J. O’Connell, Jim Deuchars, Adrian Whitehouse, Julie L. Aspden

## Abstract

LncRNAs are less conserved, yet more tissue and developmental-stage specific than mRNAs and are particularly enriched in the nervous system of *Drosophila melanogaster*, mouse and human. The function of cytoplasmic lncRNAs and their potential translation remains poorly understood. Here we performed Poly-Ribo-Seq to understand the interaction of lncRNAs with the translation machinery and the functional consequences during neuronal differentiation of SH-SH5Y cells. We discovered 237 cytoplasmic lncRNAs upregulated during early neuronal differentiation, most of which are associated with polysome complexes. The majority are cytoplasmically enriched and are intergenic or anti-sense. In addition, we find 45 small ORFs in lncRNAs to be actively translated, 17 specifically upon differentiation. 11 of these smORFs exhibit high sequence conservation across *Hominidae* suggesting they are under strong selective constraint with putative function in this clade. We discover LINC01116 is induced upon differentiation and contains an 87 codon smORF, which we detect as translated, with increased ribosome profiling signal upon differentiation. The LINC01116 peptide exhibits a cytoplasmic distribution and is detected in neurites. Knockdown of LINC01116 results in significant reduction of neurite length in differentiated cells indicating it contributes to neuronal differentiation. Our findings indicate lncRNAs are a source of non-canonical peptides and contribute to neuronal function.

## Introduction

Long non-coding RNAs (lncRNAs) are >200nt and thought to lack the potential to encode proteins. LncRNAs are well known to play key roles in development and differentiation via several mechanisms (Dimartino et al., 2018; Tsagakis et al., 2020), such as the base-pairing of lnc-31 with Rock1 mRNA, to promote its translation during myoblast differentiation (Dimartino et al., 2018). ∼40% of human lncRNAs are specifically expressed in the brain (Derrien et al., 2012), where they display precise spatiotemporal expression profiles (Ponting et al., 2009). A subset of nuclear neuronal lncRNAs have been found to regulate neuronal differentiation in mouse and human (Carelli et al., 2019; Chodroff et al., 2010; Lin et al., 2014; Winzi, 2018). Until recently, the majority of work has focused on nuclear lncRNA functions, following the belief that most lncRNAs were retained in the nucleus (Derrien et al., 2012; Djebali et al., 2012). However, it has become increasingly clear that many lncRNAs are exported to the cytoplasm (Carlevaro-Fita et al., 2016) and have specific cytoplasmic functions in post-transcriptional gene regulation whilst some possess specific neuronal roles. For example, UCHL1-AS is detected in dopaminergic neurons and promotes translation of UCHL1 mRNA by increasing its association with heavy polysomes (Carrieri et al., 2012). Whereas BACE1-AS transcript, which is significantly upregulated in the brain of Alzheimer’s disease patients, base-pairs with beta-secretase-1 (BACE1) mRNA, stabilising it by masking miR-485-5p binding sites (Faghihi et al., 2010). Alternatively, BC200 represses translation initiation in dendrites by disrupting the formation of pre-initiation 48S complexes (Wang et al., 2002).

Translation regulation is essential during neuronal differentiation (Mohammad et al., 2019), with the translation of non-canonical ORFs, e.g. upstream ORFs (uORFs), playing important roles (Blair et al., 2017; Fujii et al., 2017; Rodriguez et al., 2019). Ribosome profiling in a range of organisms and tissue types has revealed the translation of a variety of non-canonical ORFs including small ORFs (smORFs) <100 codons in length (Aspden et al., 2014; Blair et al., 2017; Duncan & Mata, 2014; Fujii et al., 2017; Guo et al., 2010; Ingolia et al., 2013; Rodriguez et al., 2019). Although these translation events remain controversial, it is clear that lncRNAs can interact with the translation machinery. Limited ribosome profiling signal found on smORFs might be the result of sporadic binding of a single ribosome and may not necessarily correspond to active translation. Poly-Ribo-Seq was previously developed to distinguish those lncRNAs that are bound by multiple ribosomes, and therefore actively translated, from such a background signal. Predicting which lncRNA-smORFs are translated has proven difficult, as attributes to predict protein-coding ability are ineffective on small non-canonical ORFs. A small but growing number of smORF peptides translated from lncRNAs have been found to exhibit cellular and organismal functions (Anderson et al., 2015; Chen et al., 2020; Magny et al., 2013; Pueyo & Couso, 2008; Spencer et al., 2020; Wang et al., 2020).

To study the dynamic interactions of lncRNAs with the translation machinery and identify actively translated cytoplasmic lncRNAs during early neuronal differentiation, we perform Poly-Ribo-Seq on SH-SY5Y cells (human neuronal cell line). During differentiation a global decrease in translation is evident and ribosomal protein mRNAs are specifically excluded from the polysomes during differentiation as part of this down-regulation. We detect ∼800-900 lncRNA genes in the cytoplasm of SH-SY5Y cells. Specifically, 237 lncRNAs are upregulated and 100 downregulated during differentiation, 58-70% of which interact with polysome complexes. These interactions are dynamic during differentiation. Ribo-Seq identified 45 actively translated smORFs in lncRNAs, several of which are regulated during differentiation. A subset of these translation events was validated by transfection tagging experiments and analysis of publicly available mass spectrometry data. High levels of sequence conservation across *Homimidae* indicates that the resulting lncRNA-smORF peptides are produced in other great apes and likely possess cellular functions. One of the lncRNA genes that we discover to be significantly upregulated during differentiation is LINC01116, which is cytoplasmically enriched and associated with polysomes. Our Ribo-Seq analysis reveals the translation of an 87aa peptide from LINC01116 with increased ribosome profiling coverage upon differentiation. The LINC01116 peptide exhibits a cytoplasmic distribution and is detected in neurites. We reveal that LINC01116 contributes to neuronal differentiation and is required for proper neurite lengthening.

## Results

### Differentiation of SH-SY5Y cells with retinoic acid results in reduced translation levels

To dissect the importance of cytoplasmic lncRNAs and their ribosome associations in early neuronal differentiation we profiled the differentiation of SH-SY5Y cells with retinoic acid (RA) for 3 days. This treatment results in neuronal differentiation as indicated by neurite elongation (Sup 1A), which can be seen by immunostaining for neuronal βIII-tubulin (TuJ1) (Fig 1A). Quantification of neurite length reveals significant elongation upon RA treatment (Fig 1B). There is also increased expression of neuronal markers; more cells express c-Fos upon differentiation (Sup 1B, C). There is a concomitant reduction in cell proliferation, as seen by reduced number of Ki67+ cells (Sup 1D, Fig 1C) as well as a reduction in the levels of pluripotency marker SOX2 (Sup 1E). These differentiated cells exhibit characteristics of outer radial glia neural progenitors and are driven towards a noradrenergic phenotype as indicated by increased Dopamine Beta Hydroxylase like Monoxygenase Protein 1 (MOXD1) expression (Sup 1F) (Pollen et al., 2015; Xin et al., 2004). Translational output assessment by polysome profiles (Fig 1D) reveals that differentiation results in down-regulation of global translation. Quantification of translation complexes across the sucrose gradients indicates that levels of polysomes are reduced with respect to 80S monosomes (Sup 1G) resulting in a reduced polysome to monosome ratio (Fig 1E). This down-regulation of translation is accompanied by a shift of ribosomal protein (RP) mRNAs from polysomes to monosomes; e.g. RpL26 mRNA (Fig 1F, Sup 1H), RpS28 (Sup 1I) and RpL37 (Sup 1J), as measured by RT-qPCR across gradient fractions. This reduced synthesis of RPs has previously been reported during neuronal differentiation (Blair et al., 2017; Chau et al., 2018).

**Figure 1:**
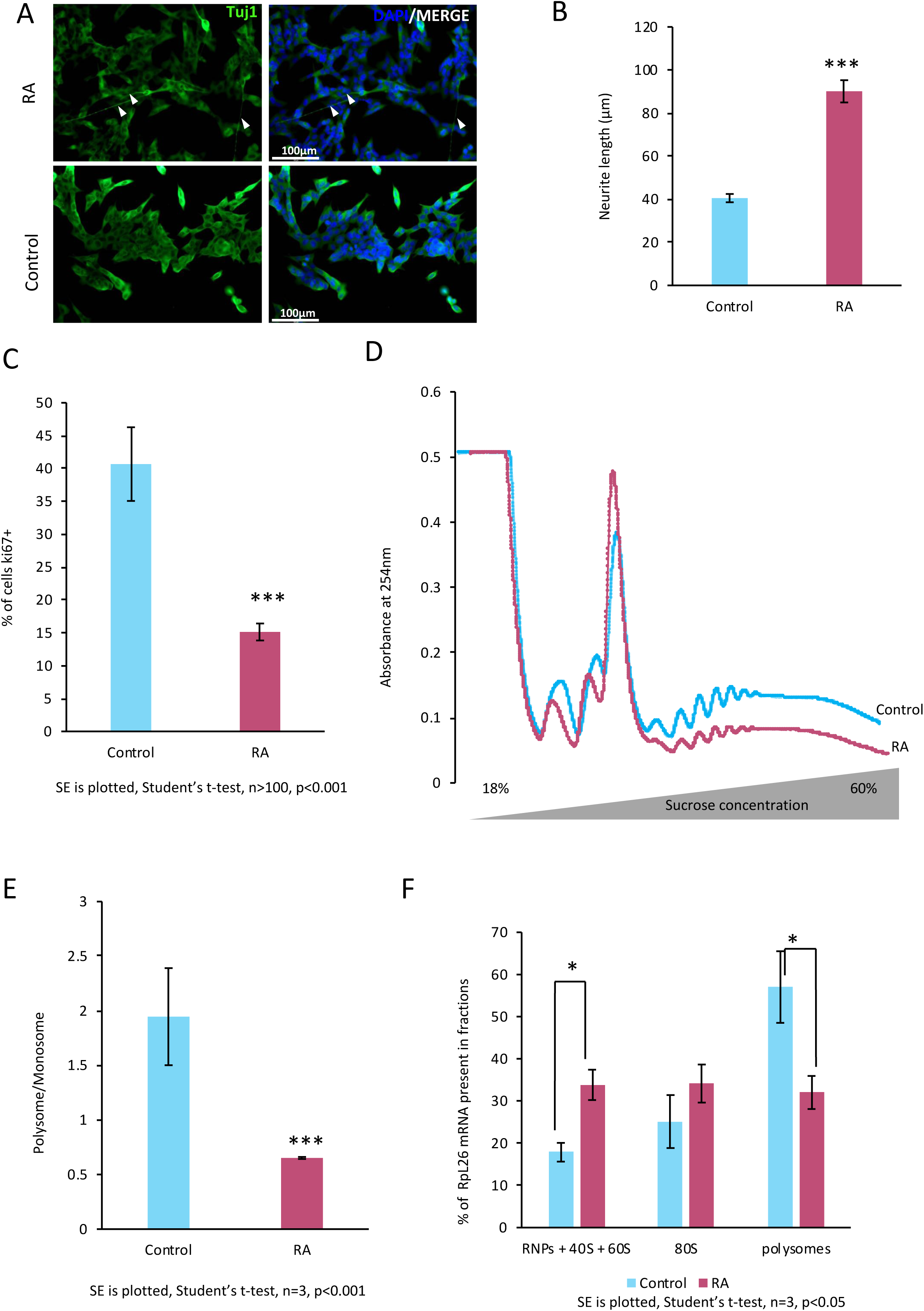
Differentiation of SH-SY5Y results in a reduction in the level of active translation. Differentiation of SH-SY5Y cells with trans-retinoic acid (RA) for 3 days results in neurite elongation as seen by (A) representative immunofluorescence after staining for TuJ1 (βlll-tubulin) (arrowheads mark elongated neurites) and (B) quantification of neurite length(Student ‘s t-test, n=3, p<0.01). (C) RA treatment also results in reduction of proliferation, indicated by the reduction of ki67+ cells (Student’s t-test, n=1000, p<0.001). (D) Sucrose gradient UV absorbance profiles of undifferentiated (Control) and differentiated (RA) cells reveals global translation is reduced upon differentiation. Peaks correspond to ribosomal subunits (40S, 60S), monosome (80S) and polysomes. (E) Quantification of Polysome/Monosome (P/M) ratio indicates reduction of active translation is significant upon RA treatment (student’s t-test, n=3, p<0.001). (F) RPL26 ribosomal protein mRNA shift from polysomes to RNPs and ribosomal subunits upon differentiation, as shown by RT-qPCR across gradient (student’s t-test, n=3, p<0.05).

### Poly-Ribo-Seq reveals differences in RNA expression and polysome association upon differentiation

To profile RNA, ribosome association and translational changes upon differentiation we employed Poly-Ribo-Seq (Aspden et al., 2014) with some minor modifications to adapt to human neuronal cells (Fig 2A). This adaptation of ribosome profiling (Ribo-Seq) can detect which RNAs are cytoplasmic, polysome associated and translated (Fig 2A). We sequenced i) poly-A selected cytoplasmic RNA, ‘Total’ RNA-Seq, ii) Polysome-associated poly-A RNAs, ‘Polysome’ RNA-Seq, and iii) ribosomal footprints, Ribo-Seq, from control and RA differentiated cells, with three biological replicates (Materials and Methods for details).

**Figure 2:**
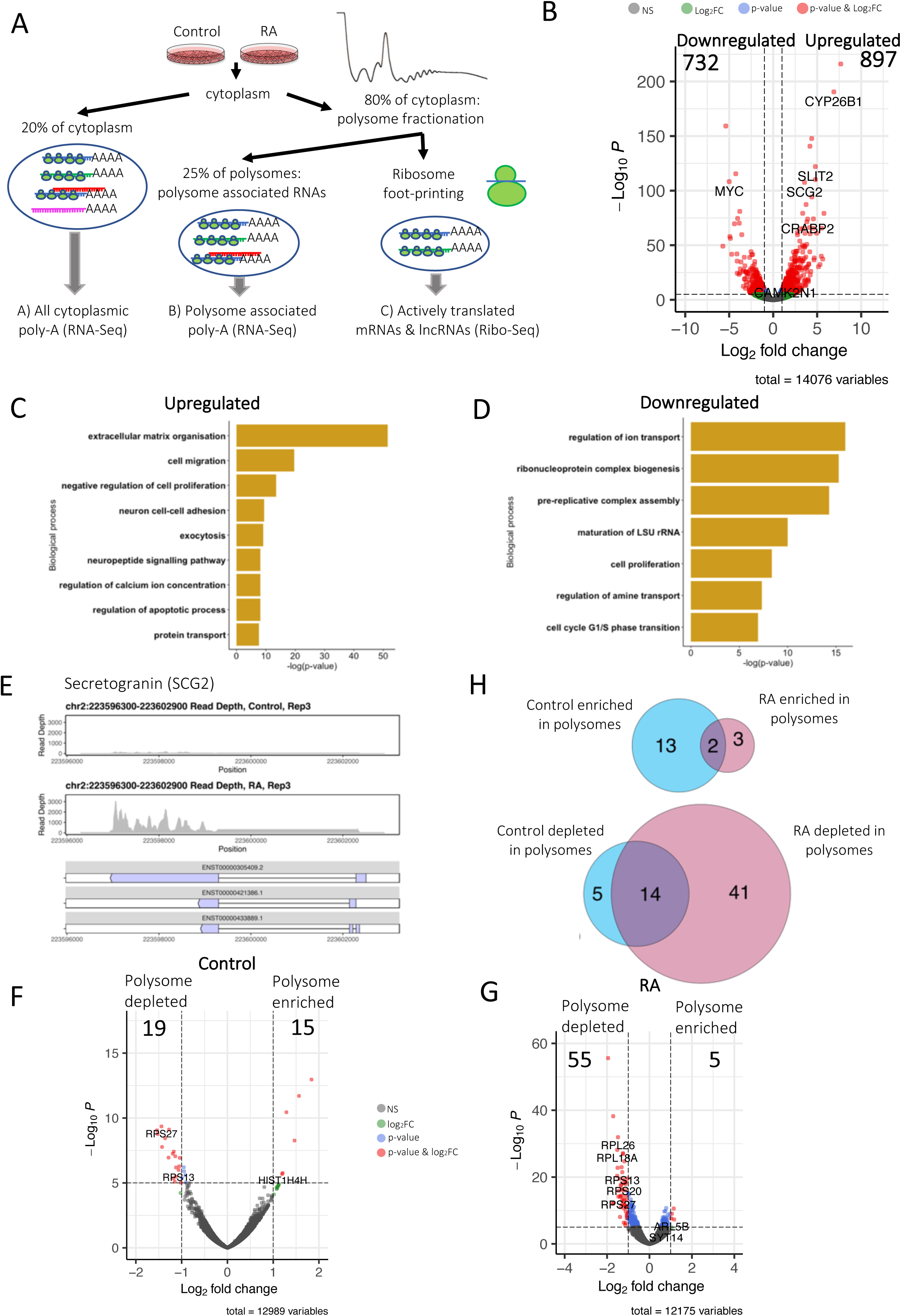
Differentiation results in global RNA changes. A) Schematic of Poly-Ribo-Seq. (B) Volcano plot of protein-coding gene level changes in response to differentiation (polysome-associated); with 897 protein-coding genes significantly upregulated and 732 significantly downregulated (log_2_ fold-change cut-off=1, p^adj^<0.05). GO terms enriched in polysome-associated protein-coding genes (C) upregulated and (D) downregulated upon differentiation. (E) Genome browser visualization of RNA-seq reads mapping to neuronal transcript Secretogranin, induced upon differentiation. (F, G) Volcano plots of differential expression of protein-coding transcripts between Total cytoplasm and polysomes for (F) Control and (G) RA, revealing the depletion of RP mRNAs from polysomes upon differentiation. (H) Venn diagrams of overlap between those mRNAs enriched or depleted in the polysomes between Control and RA.

PCA of the 3 Total and 3 Polysome RNA-Seq for both Control and RA treatment shows that RA treated samples cluster separately from Control samples and biological replicates generally cluster together (Sup 2A). To profile differences as a result of differentiation we compared Control and RA samples, performing differential expression analysis at the gene level for both Polysomal RNA-Seq samples (main figures) and Total Cytoplasmic RNA-Seq samples (supplemental figures). To dissect the ability of RNAs to associate with translation complexes we compared the “Polysomal” and “Total” RNA populations for both Control and RA conditions, again through differential expression analysis at the gene level. Differences in expression between Control and RA are greater than those seen between Total and Polysome populations. Interestingly the variation in gene expression between differentiated (RA) Total and Polysome is greater than the variation between Control Total and Control Polysome (Sup 2A).

Analysis of the two types of RNA-Seq revealed large changes in RNA expression in response to differentiation, as previously determined (Blair et al., 2017). In fact, 897 protein-coding genes are upregulated upon RA treatment and 732 downregulated, within the Polysomes (Fig 2B). A very similar pattern is seen from Total; 936 up-regulated and 691 down-regulated genes (Sup 2B), and 78% of the significantly affected protein-coding genes are in common between Total and Polysome populations (Sup 2C&D). GO term analysis of these differentiation-regulated protein-coding genes indicates that functions corresponding to neuronal differentiation are upregulated (Fig 2C). There is an enrichment of ribosome and translation functions in those genes downregulated upon neuronal differentiation (Fig 2D). A clearer enrichment of genes with neuronal functions is evident in GO analysis of Total RNA-Seq (Sup 2E), e.g. synaptic transmission and similar enrichment of translation functions are in the downregulated set (Sup 2F). Expression of genes required for neuronal differentiation is increased or induced upon RA treatment, e.g. Secretogranin, which is involved in packaging of neuropeptides into secretory vesicles (Sossin & Scheller, 1991) (Fig 2E).

Comparison of Total and Polysome RNA-Seq indicates that some mRNAs are enriched in polysomes whilst others are depleted. Ribosomal protein mRNAs are specifically depleted from polysomes upon differentiation, as seen by their altered relative levels between Total and Polysomes in Control (Fig 2F) and RA (Fig 2G). This is consistent with a global downregulation in translation (Fig 1D) and shift of RP mRNAs out of the polysomes (Fig 1F). By comparing these polysome enriched and depleted mRNAs between Control and RA the dynamic nature of polysome association during differentiation is revealed (Fig 2H). More mRNAs become depleted during differentiation, including more RP mRNAs. GO term analysis of those genes enriched or depleted in the polysomes (Sup 2G, H, I) supports this depletion of RP mRNAs upon differentiation (Sup 2I).

### LncRNAs are induced and associated with polysomes during differentiation

To understand the potential role and regulation of cytoplasmic lncRNAs we analysed their expression levels in Total and Polysome populations upon differentiation (Sup 3A). We detect large numbers of lncRNAs present in the cytoplasm. 801 lncRNA genes expressed and present (i.e. have RPKMs ≥ 1) in the cytoplasm under control conditions and 916 lncRNA genes in differentiated cells. Interestingly, 237 lncRNA genes were upregulated during differentiation whilst only 82 were downregulated, within Polysome fractions (Fig 3A). A very similar pattern was seen when considering Total RNA-seq with 178 lncRNA genes up-regulated and 100 down-regulated (Sup 3B). The majority of these differences were identified in both Total and Polysome populations (70% of the upregulated and 58% of the downregulated lncRNAs were in common between total and polysome) (Sup 3C&D). Interestingly when we look in more detail at these lncRNA regulated during differentiation the majority of differentiation-regulated lncRNAs are either intergenic or/and anti-sense lncRNAs (Fig 3 B&C-Polysome, Sup 3 E&F-Total). Direct comparison of levels of lncRNAs within the whole cytoplasm and within polysome fractions indicates the vast majority lncRNAs (Control: 98% and RA: 99% are neither polysome enriched nor depleted (Sup 4 A&B). A small number (Control: 12 and RA: 10) of lncRNAs are specifically depleted from the polysomes including several lncRNAs derived from small nucleolar RNA host genes. Interestingly there is a smaller proportion of anti-sense lncRNAs within these polysome-depleted populations (Sup 4 C&D) compared to differences in expression during differentiation (Fig 3 B&C).

**Figure 3:**
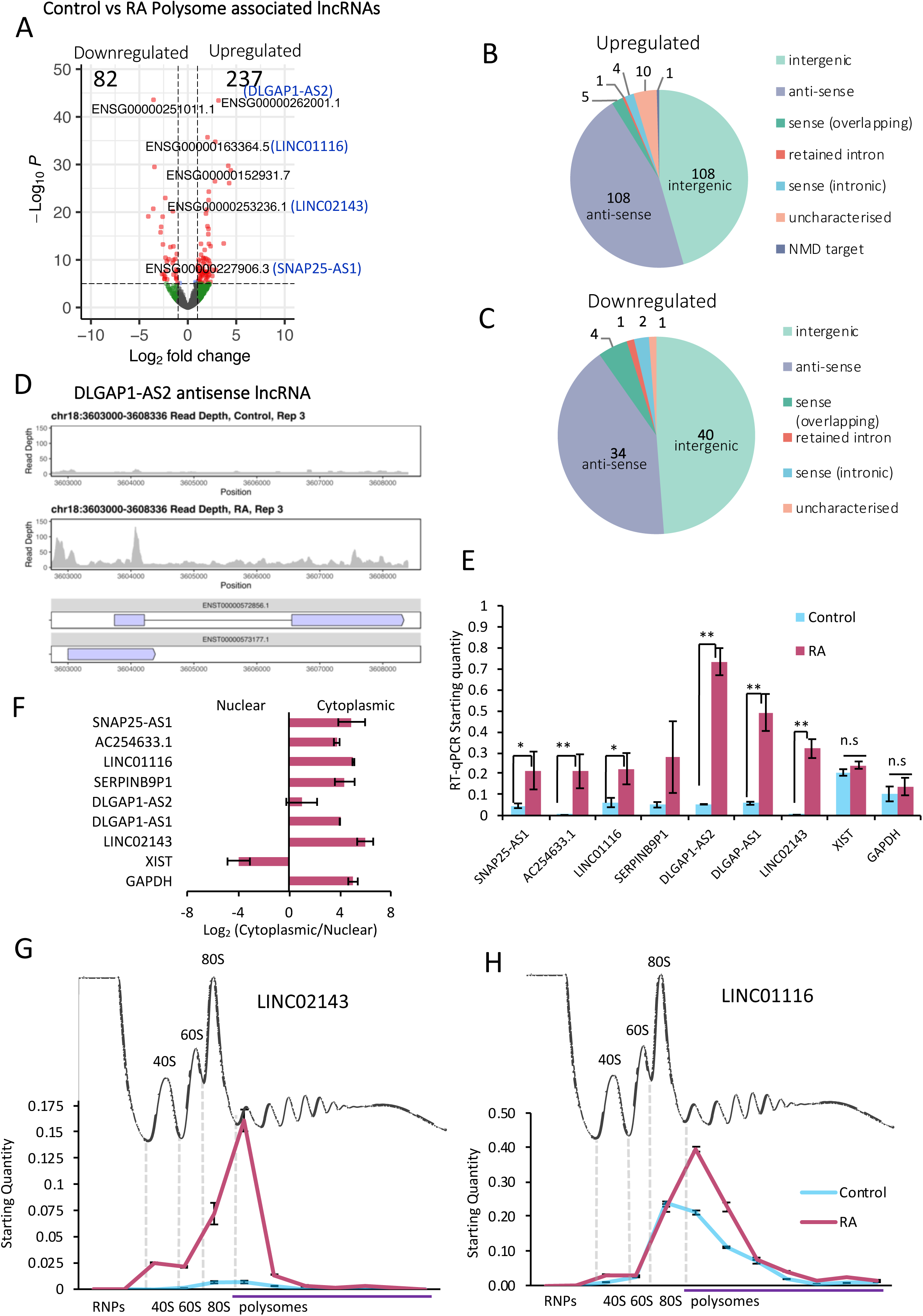
LncRNAs regulated and associated with polysomes. (A) Volcano plot displaying the differentially expressed polysome-associated lncRNAs (labelled by their geneIDs) between Control and RA populations. 237 lncRNAs are upregulated during differentiation and 82 downregulated (log_2_ fold-change cut of f=1, p^adj^<0.05). Pie charts showing types of lncRNAs (B) upregulated and (C) downregulated upon differentiation (intergenic; anti-sense; sense-overlapping; retained intron; sense-intronic; uncharacterized; NMD target). (D) Example of lncRNA induced upon differentiation, DLGAP1-AS2. RNA-seq coverage for DLGAP1-AS2 in Control and RA conditions. (E) Changes in lncRNA levels upon differentiation validated by RT-qPCR, with XIST lncRNA and GAPDH mRNA as controls (SE is plotted, n=3, student’s t-test p<0.05 and p<0.01). (F) LncRNAs of interest that are induced are specifically localised to cytoplasm as shown by subcellular fractionation RT-qPCR. XIST lncRNA was used as a nuclear and GAPDH mRNA as a cytoplasmic positive control (n=3, SE is plotted, student’s t test, n=3, p>0.05). RT-qPCR of IncRNAs across sucrose gradient fractions indicates that (G) LINC02143 is found in 80S and small polysome fractions during differentiation (in differentiated cells 5% of the transcripts is detected in 80S (monosome) fraction and 65.6% in small polysome complexes) and (H) LINC01116 is found in 80S and 2-7 polysome fractions both in control and RA treated cells. On average, 66% of the LINC01116 transcripts is detected in the polysome fractions in Control and 57% upon differentiation. (n=3, SE is plotted).

Significant induction of specific polysome-associated lncRNAs during differentiation, such as DLGAP1-AS2 is suggestive of a regulatory role during neuronal differentiation (Fig 3 A&D). We validated these differentiation-induced changes in a subset of 7 lncRNAs (Fig 2E). By RT-qPCR the expression of all candidate lncRNAs were significantly upregulated upon differentiation, as was determined by RNA-Seq analysis. Fold-changes were highly correlative between RNA-Seq and RT-qPCR (Sup 4E). To enable us to focus on lncRNAs with potential neuronal functions we selected candidate lncRNAs that exhibited the highest fold increase in levels upon differentiation. The majority of these candidate lncRNAs (6/7) are specifically enriched in the cytoplasm, rather than the nucleus (Fig 3E, Sup 4F) in contrast to the known nuclear lncRNA Xist. Only 2/7 lncRNAs tested showed changes in cytoplasmic enrichment upon differentiation (Sup 4G), the majority do not (5/7). When these candidate lncRNAs were profiled across sucrose gradient fractions they were found to associate with polysome complexes within the cytoplasm. LINC02143, which is induced >22-fold during differentiation (Fig 3E), is highly enriched in the cytoplasm compared to the nucleus (Fig 3F) and is found in monosomes and small polysomes (Fig 3G). It is also just as upregulated in polysomes (Fig 3A) as in the total cytoplasm (Sup 3B) during differentiation. Unlike DLGAP1-AS2, DLGAP1-AS1 is enriched in the cytoplasm (Fig 3F) and associates with small polysomes, as well as ribosomal subunits (Sup 4H).

Another lncRNA whose levels significantly increase during differentiation is LINC01116 (Fig 3 A&E), which is involved in the progression of glioblastoma (GBM) (Brodie et al., 2017). LINC01116 is enriched in the cytoplasm (Fig 3F), detected at high levels in the 80S (monosome) fraction and in small polysomal complexes (Fig 3H). Upon differentiation there is an increase in the amount of LINC01116 present in disomes, compared to Control. This is consistent with the upregulation of LINC01116 transcript in the polysomes detected by RNA-Seq, indicating a functional interaction of LINC01116 with polysomes during differentiation. In fact, the majority of LINC01116 transcript was found to associate with polysomes in both undifferentiated (Control) cells (66%) and upon differentiation (RA-57%), suggesting it is could either be translated or associating with translation complexes (Fig 3H).

### Translation of lncRNA-smORFs during differentiation

To better understand the association of lncRNA with polysome complexes and their potential translation we analysed ribosome footprinting from our Poly-Ribo-Seq experiments (Fig 2A). Framing analysis reveals good framing, specifically at footprint lengths of 31 and 33 nt (Fig 4A) and discrete footprint lengths (Sup 5A). On average 95% of ribosome footprints mapped to CDSs, whilst in RNA-Seq this was only 53% (Sup 5B). Metagene analysis of 31 nt length reads indicates that framing is high within ORFs, and that the footprint signal outside of ORFs is mostly within 5’-UTRs rather than 3’-UTRs, where there is a sharp drop off in footprints at the stop codon (Sup 5C). Together these attributes indicate that our Ribo-Seq quality is high and represents genuine translation events. Since 31 and 33nt reads exhibited high triplet-periodicity they were selected for downstream analysis of translation events to identify ORFs that are translated (Fig 4B). Overall, we are able to detect 16,282 protein-coding ORFs translated in Control and 12,745 in RA, 10,014 of which are translated in both conditions (Table 1). Interestingly we are also able to detect the translation of 71 uORFs and 5 dORFs (Table 1).

**Table 1:**
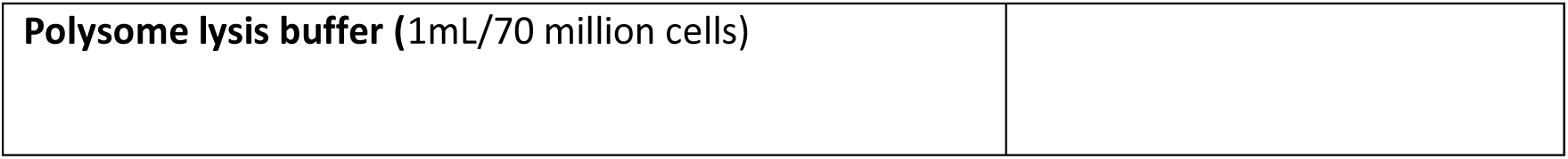
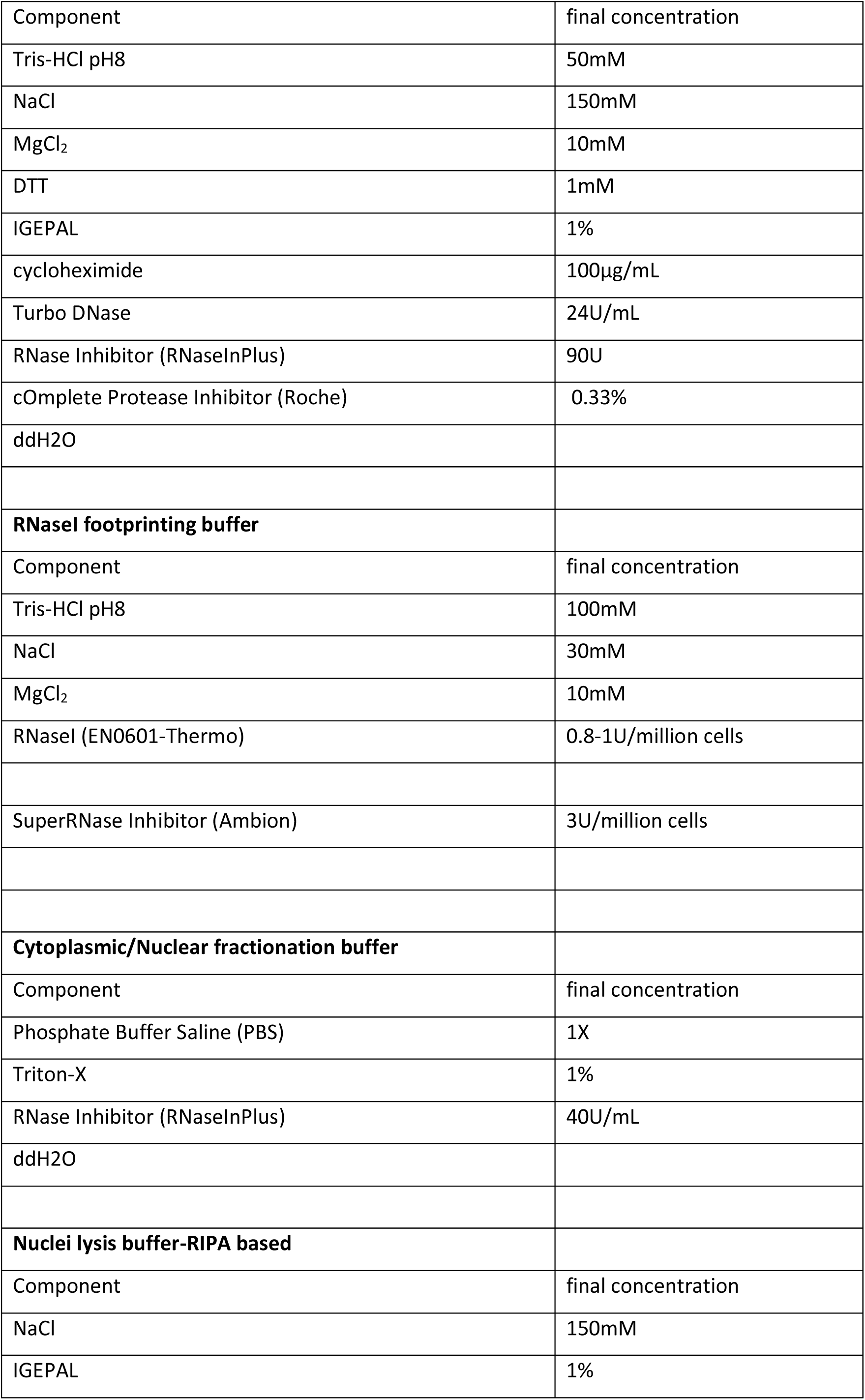
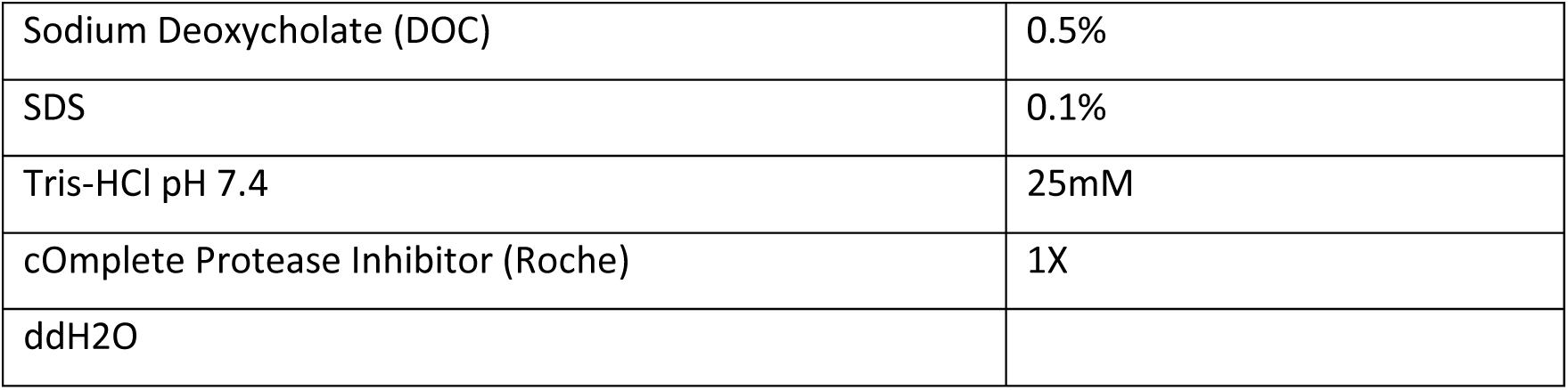
Buffers.

**Table 2:**
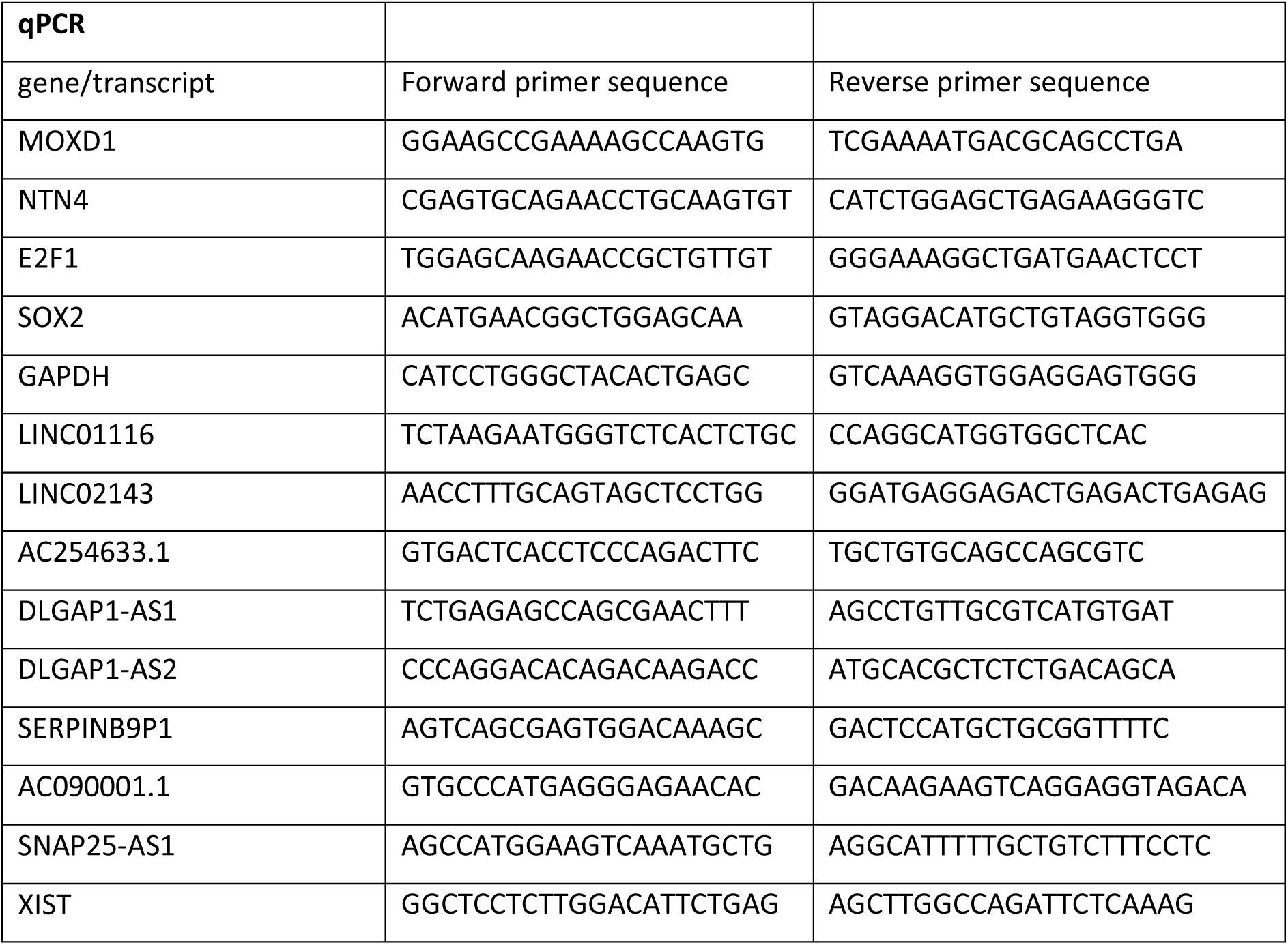
qPCR primers.

**Figure 4:**
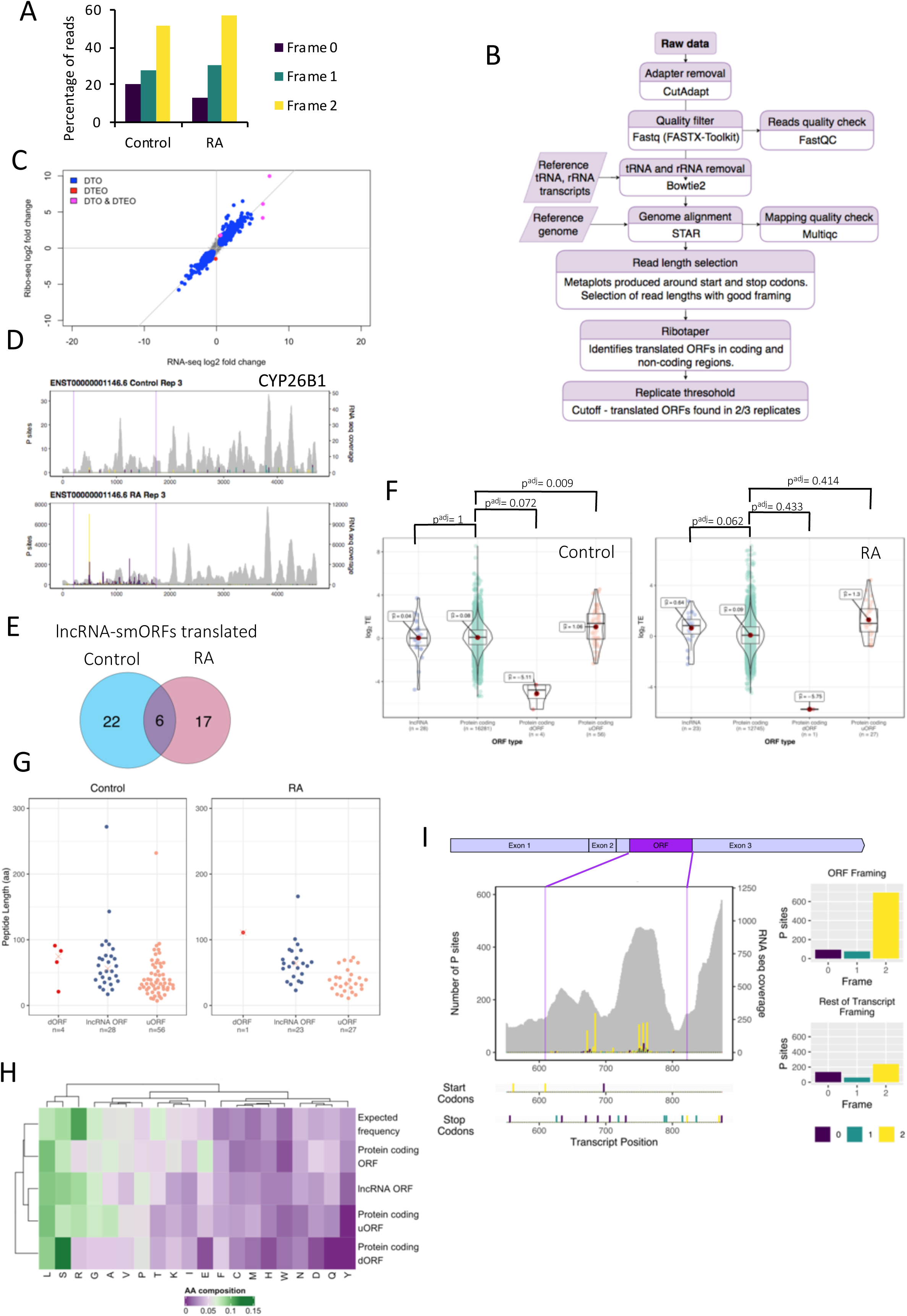
Translation of lncRNA smORFs. (A) Poly-Ribo-Seq exhibits triplet periodicity, reflecting genuine translation events. (B) Workflow for identification of translated ORFs from ribo-seq and RNA-seq, see materials and methods for details.(C) Plot of log2-fold change for each ORF in RNA (RNA-Seq) versus ribosome footprints (Ribo-Seq). DTOs (differentially transcribed ORFs) or forwarded ORFs are driven by transcriptional regulation, with significant ΔRPF and ΔRNA but no significant ΔTE (blue). DTEOs (differential translation efficiency ORFs) or exclusive ORFs are driven by translational regulation, with significant ΔRPF and ΔTE without any significant ΔRNA (red). DTO & DTEOs (ORFs undergoing differential transciption and translation efficiency) are either intensified; a significant ΔTE acts with a significant ΔRNA, or buffered; a significant ΔTE is completely counteracting a ΔRNA, leading to no significant ΔRNA(pink). (D) View of Poly-Ribo-Seq data for CYP26B1. This is DTG ORF and exhibits significant ΔRNA and ΔRPF. (E) Venn diagram of lncRNAs smORFs translated in Control and RA, with overlap. (F) Plots of Translational Efficiencies for protein-coding ORFs, lncRNAs-smORFs, dORFS and uORFs. (G) Length distribution of translated ORFs in lncRAs, dORFs and uORFs (in codons) in Control and RA. (H) Amino acid usage for smOFs compared to protein-coding ORFs and expected frequency. (I) Poly-Ribo-Seq profile for LINC01116 in RA treatment. RNA-Seq (Polysome) reads are grey and ribosome P sites in purple, turquoise and yellow according to frame. Purple lines mark beginning and end of translated smORF. All possible start and stop codons are indicated below. Framing within and outside translatd smORF shown on left.

To monitor translational changes we performed differential translation analysis (Chothani et al., 2019). Differential ribosome occupancy on ORFs (ΔRPF), differential mRNA expression (ΔRNA) and differential translation efficiency (ΔTE) were calculated (Chothani et al., 2019). The majority of changes in protein production are driven by transcriptional regulation during differentiation, with no change in TE (Fig 4C). We detected 3 protein-coding CDSs being exclusively regulated by translation, upon differentiation (Fig 4C). One of these is centromere protein S (CENPS), which is expressed at low levels in neuroblastoma tumours and is thought to reduce cell growth (Carén et al., 2005; Krona et al., 2004). We were also able to identify CDSs that are known to be regulated during differentiation such as cytochrome P450 26A1 and cytochrome P450 26B1 (CYP26A1, CYP26B1), involved in RA metabolism (Maden, 2007) (Fig 4D). These ORFs are transcriptionally and translationally upregulated, and their TE is significantly increased upon differentiation.

Importantly, using our pipeline (Fig 4B) we also detect the translation of 45 ORFs within lncRNAs, 28 in control and 23 during differentiation, 6 of these were translated in both conditions (Fig 4E). Interestingly, TEs for these lncRNA-smORFs are similar to those of protein-coding ORFs (Fig 4F), providing further evidence that these are genuine translation events. The dORFs that we detect are translated at much lower efficiencies (Fig 4F). This makes sense considering that ribosomes will likely have to reinitiate after the main ORF, which will occur at a low efficiency. One of the lncRNA-smORFs we identify to be translated in differentiated cells, has previously been characterised, the 84 aa CRNDEP (Szafron et al., 2015). Analysis of these 45 translated smORFs from lncRNAs shows that they are <300 aa in length with a median size of 56 aa in Control and 64 aa in RA (Fig 4G, Sup 5D).

Previous analysis has indicated that smORF peptides exhibit specific amino acid usage, which suggests they are both genuine proteins and show enrichment for transmembrane alpha-helices (Song et al., 2010). Therefore, we profiled the amino acid composition of our lncRNAs-smORFs, uORFs and dORFs compared to protein-coding ORFs and expected frequency (Fig 4H). lncRNA-smORFs exhibit similar frequencies to known protein-coding ORFs. Specifically, smORFs possess lower than expected arginine levels, but not as low as known protein-coding ORFs. Amino acid usage does not suggest that any smORF groups have propensity to form transmembrane alpha-helices (Aspden et al., 2014).

From within the set of lncRNAs we identified as induced during differentiation, several contained translated smORFs. One of these is LINC01116, whose translated ORF exhibits high framing (Fig 4I). In fact, ∼80% of reads that map to this ORF are in frame 2, whereas outside this ORF, the few reads mapping to the remaining lncRNA sequence are far more equally distributed between the three possible frames (Fig 4I). This smORF is 71-codons, and its start codon is 608nt into the lncRNA. Ribosome profiling signal is substantially higher upon differentiation, mainly as a result of increased lncRNA transcript abundance (Sup 5E).

### Peptide synthesis from smORFs in lncRNAs during differentiation

Our pipeline is highly stringent, i.e., there are many additional ORFs that display good framing but do not pass our thresholds for numbers of footprinting reads or exhibit background signal in rest of the lncRNA. Therefore, we are confident these translation events are taking place. To validate peptide synthesis, we have taken two complementary strategies; mass spectrometry analysis and transfection of ORF tagging constructs. Mass spectrometry generally only detects 90% of translated proteins from protein-coding genes (Zubarev, 2013). Analysis of previously published mass spectrometry datasets from SH-SY5Y (undifferentiated and RA-treated) cells (Brenig et al., 2020; Murillo et al., 2018) supports the production of peptides from 6 of our translated uORFs (2 Control, 4 RA) and 8 lncRNA-smORFs (4 Control and 4 RA) (Fig 5A). This represents ∼8% (uORFs) and 18% (lncRNA-smORFs) of what is detected in Poly-Ribo-Seq. This is to be expected given the small size of these peptides and therefore reduced chance of producing peptides >8aa from digestion for mass spectrometry detection (Saghatelian & Couso, 2015).

**Figure 5:**
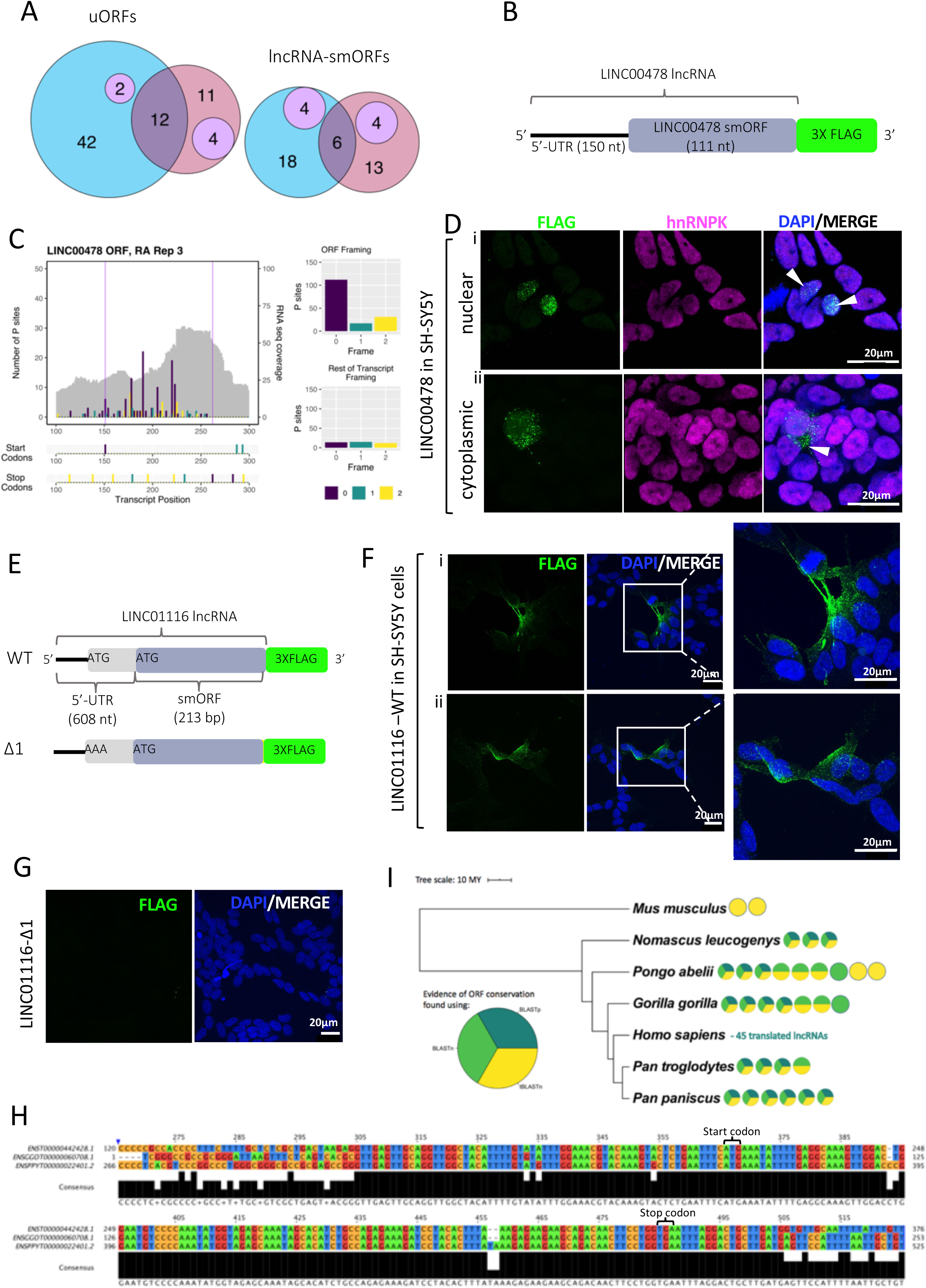
Peptide production from smORFs in lncRNAs. (A)Venn diagrams showing overlap in uORFs and lncRNA-smORFs detected, between our Poly-Ribo-Seq and publicallyav ailable mass spectrometry data from SH-SY5Y (purple). Control in blue, RA in pink. (B) Schematic of tagging construct for LINC00478; lncRNA sequence upstream of smORF and smORF, excluding its stop codon, cloned upstream of 3X FLAG, which is lacking its own start codon. FLAG signal is therefore dependent on smORF translation. (C) Poly-Ribo-Seq profile for LINC00478 in RA treatment. RNA-Seq (Polysome) reads are grey and ribosome P sites in purple, turquoiseand yellow according to frame. Purple lines mark beginning and end of translated smORF. Allpossible start and stop codons are indicated below. Framing withinand outside translated smORF shown on right. (D) Confocal images of FLAG-tagged LINC00478 peptide in SH-SY5Y cells (Control), showing (i) nuclear and (ii) cytoplasmic distribution, green is FLAG, magenta is hnRNPK (marking nuclei) and blue is DAPI (scale bar is 20μm). (E) Schematic of tagging constructs for LINC01116 (WT and start codon mutant Δ1). (F) Confocal images of FLAG-tagged LINC01116 peptide (WT showing cytoplasmic localisation, near cell membrane and neuritic processes (magnification of insert is 3X). (G) Δ1 start codon mutant, showing no FLAG signal, in SH-SY5Y cells; green is FLAG and blue is DAPI (scale bar is 20μm). (H) Portion of human ENST00000442428 lncRNA nt alignment with gorilla and orangutan ntsequences, showing the smORF. Alignment built in ClustalOmega (Sievers, F. et al. 2011) displayed in JalView (Waterhouse, A.M. et al. 2009). (I) Phylogram with lncRNA-smORFs for which evidence of sequence conservation were found represented as circles, coloured according to how sequence conservation was identified. Phylogram built in iTOL (Letunic, I. et al .2006) using data from TimeTree (Kumar, S., et al. 2017), scale in 10 MYA along.

To validate translation of our lncRNA-smORFs that were not identified in previous mass spectrometry but were detected as translated by our Poly-Ribo-Seq analyses, we have also used a transfection tagging approach. We cloned the lncRNA sequence to the 5’ of the putative ORF, termed the 5’-UTR and the smORF itself, without its stop codon, with a C terminal 3x FLAG tag (Fig 5B). The start codon of this FLAG tag is deleted, so any FLAG signal is the result of translation from the lncRNA-smORF. Two candidate lncRNA-smORFs were selected that did not have mass spectrometry support; LINC000478 and LINC01116. A 37 codon smORF was detected as translated from LINC00478 in both conditions (Fig 5C). Transfection of LINC00478-smORF-FLAG into undifferentiated SH-SY5Y cells produced FLAG signal in both nuclear and cytoplasmic compartments (Fig 5D & Sup 6A). FLAG signal was also seen when LINC00478-smORF-FLAG transfected SH-SY5Y cells were treated with RA (Sup 6A). Interestingly this RA FLAG signal was only ever detected in the nucleus. Similar results were seen in HEK293 cells (Sup 6B), but because of the higher transfectional efficiency in HEK293 compared with SH-SY5Y cells, we detect FLAG signal in more cells.

Poly-Ribo-Seq detected increased levels of LINC01116 during differentiation (4-fold in Control vs RA polysome) and translation of a 71-codon smORF during differentiation (Fig 4I). Tagging of LINC01116-smORF (Fig 5E) generated FLAG signal in the cytoplasm of SH-SY5Y cells, which is localized to neurites (Fig 5F), suggesting it could play a role in differentiation. FLAG signal was also present in LINC01116 transfections in HEK293 cells (Sup 6C), but because of the higher transfectional efficiency in HEK293 compared with SH-SY5Y cells, we detect FLAG signal in more cells.

LINC01116-smORF has two potential ATG start codons (Fig 5E and Sup 6D). When these start codons were assessed for similarity to the Kozak sequence consensus, using NetStart1.0 (Pedersen & Nielsen, 1997), both exhibited scores >0.5, indicating both are in good context and therefore either could be used to initiate translation (AUG_1_= 0.545, AUG_2_= 0.645). Given the scanning model of translation initiation it seems likely that the first AUG would be used. To determine if the first start codon is actually used it was mutated (Fig 5E). No FLAG signal was present in transfections where the 5’ start codon was mutated (Δ1) (Fig 5G). This suggests that the first start codon is necessary for the translation of the LINC01116-smORF and the resulting peptide is 87aa long. Although FLAG signal is present in a low number of cells, no transfection controls and Δ1 indicate that FLAG signal is dependent on translation of the LINC01116-smORF (Sup 6E).

### Translated lncRNA-smORFs exhibit sequence conservation

Another indicator of coding potential and also of peptide functionality is the sequence conservation of our translated lncRNA-smORFs. To ensure detection of sequence conservation for these short smORFs irrespective of annotation in other genomes, we used three complementary BLAST strategies using transcript nt sequence, smORF nt sequence and protein aa sequence.

Initial searches using the entire lncRNA transcript sequence (nt) and BLASTn (Altschul et al., 1990), returned results for ∼78% of translated lncRNAs, many of which had short alignment lengths of 30-100 nt. Although some of these results may represent conservation of the smORFs, many are due to small areas of sequence overlap along the rest of the lncRNA. This is a well-documented issue when investigating lncRNAs; they rarely exhibit the same levels of conservation as mRNAs (Johnsson et al., 2014). A more common finding is that there are “modules” of higher sequence conservation across lncRNA transcripts, as described for Xist lncRNA (Brockdorff, 2018). To accommodate this, a second round of searches were performed, on the initial search results, using the nt sequence of the smORF (BLASTn) (Altschul et al., 1990). This filtering step resulted in fewer, higher quality results, with 14 of 18 remaining lncRNA-smORFs passing manual cross validation. For the majority of the these lncRNA-smORFs, conservation is high across the body of the smORF, and drops off across the rest of the transcript. The ENST00000442428.1 (AL162386.2)-smORF exhibits high sequence conservation when compared to gorilla and orangutan (*Pongo abelii*), with 100% and 99% ORF nt sequence identity respectively (Figure 5H). When entire transcripts are aligned, this percentage sequence identity drops to 74% with gorilla (ENSGGOT00000060708.1), and 65% with orangutan (ENSPPYT00000022401.2), indicating the ORF is the most conserved part of these transcripts.

To further corroborate these results, a tBLASTn (Altschul et al., 1990) search of the lncRNA-smORF aa sequences was performed. By translating transcript databases into aa in all 6 frames, tBLASTn removes the noise of synonymous substitutions, which can have a significant effect, particularly in smORFs (Ladoukakis et al., 2011). For the majority of smORFs, the same results were returned, and evidence of conservation was found for a further three lncRNA-smORFs (ENST00000454935.1_477_633, ENST00000557660.5_42_186, ENST00000453910.5_151_262) that appear to have undergone some frameshift mutations.

At the aa sequence level, ∼18% of the translated lncRNA-smORFs returned homologous proteins from BLASTp searches (Altschul et al., 1990). All the returned proteins were unreviewed, uncharacterised proteins, with no evidence at protein, transcript or homology levels in the Uniprot database (Bateman et al., 2019). This potentially suggests that automated annotation pipelines recognised the coding potential of these smORFs, unlike in the more curated human genome annotation. Overall, the combination of these 3 layers of sequence conservation analysis reveal that 11 of our translated lncRNA-smORFs exhibit sequence conservation across *Hominidae*, 3 addition smORFs are also found in gibbons (Fig 5I), with evidence for 2 translated smORFs detectable at the greater evolutionary distance of human to mouse, but not found in all apes (Sup 6E).

### LINC01116 contributes to neuronal differentiation

To dissect the potential role of LINC01116 and its translation during differentiation we knocked down LINC01116 using a siRNA pool in both undifferentiated and differentiated SH-SY5Y cells (Fig 6A). LINC01116 knockdown initially had a limited effect on cell viability, which recovered to no effect after 2 days (Sup 7A). Interestingly, LINC01116 knockdown resulted in a significant reduction of neurite length in RA treated SH-SY5Y cells (Fig 6B, zoom in Fig 6C), compared to scrambled siRNA treated SH-SY5Y (Fig 6D). There was no effect of the knockdown in undifferentiated cells (Fig 6D). This suggests that LINC01116 is involved in the regulation of neuritic processes formation. To examine potential effects of LINC01116 knockdown further on differentiation we assessed the expression levels of the noradrenergic marker MOXD1, which is important in in neural crest development. LINC01116 siRNA knockdown, upon differentiation resulted in a reduction of MOXD1 expression levels, further indicating a role of LINC01116 in neuronal differentiation (Fig 6E). However, LINC0116 knockdown had no effect on proliferation, as measure by % of Ki67+ cells (Sup 7B & Fig 6F) or cell cycle, as measured by E2F1 mRNA RT-qPCR (Fig 6G). LINC01116 likely functions early in the differentiation pathway since its levels are significantly upregulated within the first 24 hours of RA induced differentiation (Sup 7C). Expression of LINC01116 then declines rapidly by day 8 (Fig 7D). Together these results suggest that LINC01116 functions during early differentiation, contributing to neurite process formation.

**Figure 6:**
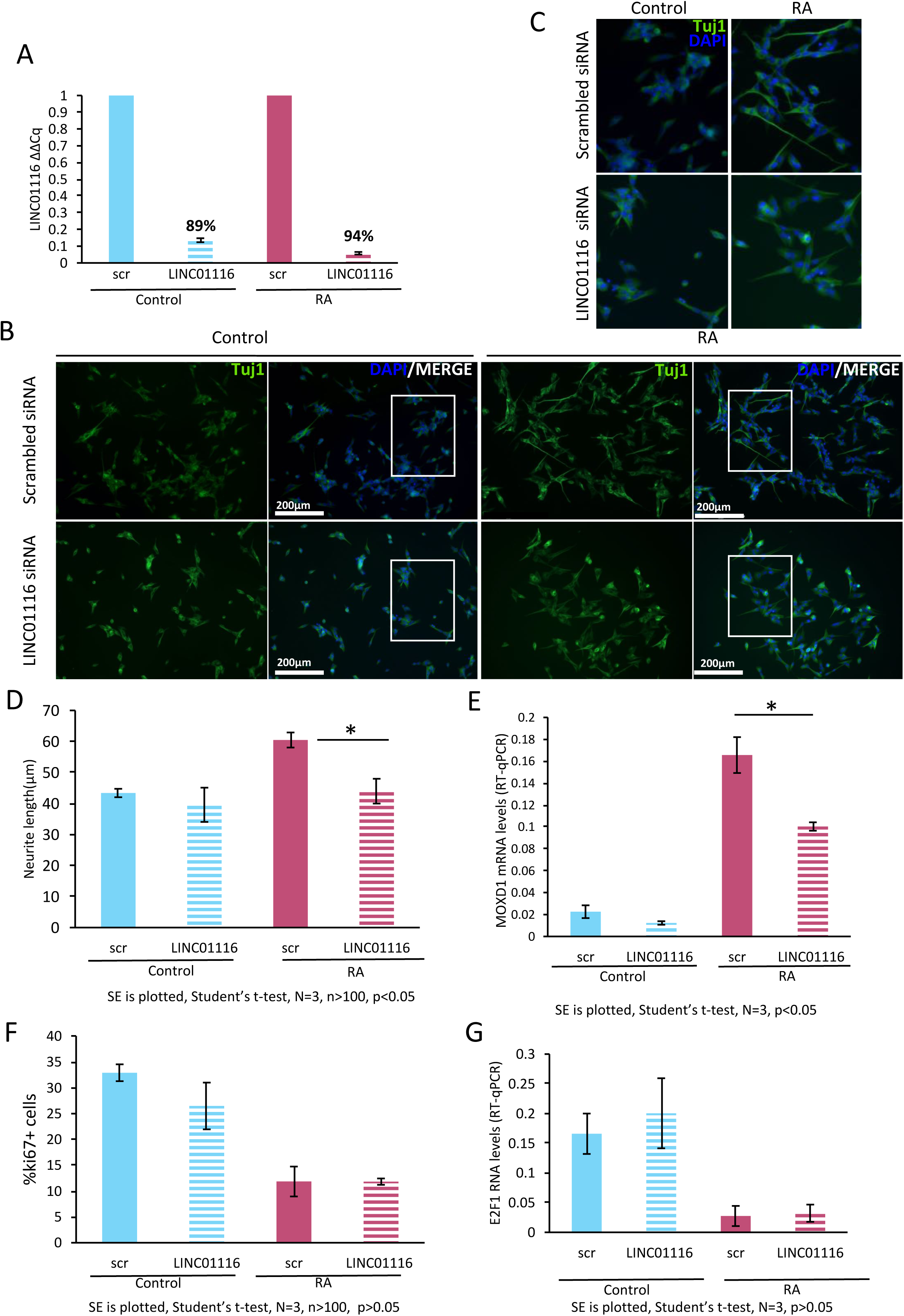
LINC01116 contributes to neuronal differentiation but does not affect cell cycle progression. (A)LINC01116 was efficiently knocked down prior to differentiation and the knockdown is persistent throughout differentiation. (B) Representative immunofluorescence image of Control and RASH-SY5Y cells, transfected with siRNA targeting LINC01116 and scrambled control,afters taining for Tuj (βIII-tubulin)at day 3 post-differentiation (scale bar=200μm). White windows magnified in (E). (C) Quantification of neurite length in Control and RA treated cellsupon knockdownshows asignificant reductionof neurite length in the differentiated cells that lack expression of LINC01116 (N=3 biological replicates, n>100 measurements, student’s t-test p<0.05). (D) RT-qPCR of differentiation marker MOXD1 in Control and RA treated cells, transfected with siRNA targeting LINC01116 and scrambled control, shows significant reduction of MOXD1 expression in differentiated cells with reduced LINC01116 levels at day 3 post-differentiation (n=3 biological replicates, student’s t-test p<0.05). (F) Quantification of proliferating cells (% of ki67+ cells) in Control and RA cells shows no effect of LINC01116 knockdown on cell proliferation (N=3 biological replicates, n>100 measurements, student’s t-test p>0.05). (G) LINC01116 knockdown does not affect cell cycle progression as shown by RT-qPCR targeting cell cycle promoting transcription factor E2F1(n=3 biological replicates, student’s t-test p>0.05).

## Discussion

Cytoplasmic lncRNAs have recently emerged as important regulators of several cellular processes, including the regulation of translation, in a variety of tissues (Carrieri et al., 2012; Dimartino et al., 2018). Moreover, the active translation of smORFs from within lncRNAs has been characterized in a range of organisms (Anderson et al., 2015; Aspden et al., 2014; Chen et al., 2020; Magny et al., 2013; Pueyo & Couso, 2008; Wang et al., 2020). Translational regulation is a key process during neuronal differentiation (Blair et al., 2017; Chau et al., 2018), therefore we reasoned that lncRNAs may functionally interact with polysomes during differentiation.

Having established that early neuronal differentiation of human neuroblastoma cells (SH-SY5Y) results in reduced global translation, we sought to further investigate the relationship of lncRNAs with the translation machinery during differentiation. We detected the presence of ∼800-900 lncRNA genes expressed and their transcripts present in the cytoplasm. 85-90% of these cytoplasmic lncRNAs are associated with polysome complexes, suggesting that they are either being translated, or regulating the translation of the mRNAs with which they interact. Moreover, the association of lncRNAs with polysomes is dynamic during differentiation, as shown by the differential polysome enrichment of lncRNAs in Control and RA treated cells. The lncRNAs that dynamically interact with polysomes upon neuronal differentiation mainly belong to the ‘intergenic’ and ‘anti-sense’ categories of lncRNAs. These results reveal that many lncRNAs are present in the cytoplasm, enriched there, and are associated with translation complexes.

An example of a polysome-associated lncRNA that we characterized in more detail is LINC02143. It is an intergenic lncRNA with no known function, which we find induced upon differentiation. It is detected in 80S and small polysome fractions, indicating it interacts with the translation machinery. A number of anti-sense polysome-associated lncRNAs appear to be upregulated upon differentiation. Amongst them is DLGAP1-AS1, which is anti-sense to Disks large-associated protein 1 (DLGAP1), itself involved in chemical synaptic transmission. DLGAP1-AS1 interacts with actively translating polysomes both in Control and upon differentiation. The lncRNAs depleted from the polysomes have fewer anti-sense lncRNAs relative to other populations, suggesting that anti-sense lncRNAs are preferentially localised to polysomes. These polysome associated anti-sense lncRNAs could potentially regulate the translation of their ‘sense’ mRNA, through base-pairing, as is the case with BACE1-AS (Faghihi et al., 2010) and UCHL1-AS (Carrieri et al., 2012). This contrasts with the general view that anti-sense lncRNAs regulate gene expression in the nucleus at the point of transcription or splicing (Beckedorff et al., 2013).

Differential translation analysis revealed that upon RA induced neuronal differentiation of SH-SY5Y for 3 days, a small number of ORFs are exclusively translationally regulated. For the majority of ORFs, translation regulation is driven by transcription regulation. This result indicates that at this early timepoint, transcriptional control, rather than translational control is the driving force of SH-SY5Y differentiation. This is not surprising, because according to a previous study of human embryonic stem cells (hESCs) differentiation into neural progenitor cells (NPCs) only 12% of the detected transcripts were exclusively translationally regulated (Blair et al., 2017).

Ribosome profiling of the actively translating polysomes allowed us to distinguish between the lncRNAs that associate with the polysome complexes and those that are being actively translated. We were able to identify 45 translated lncRNA-smORFs in undifferentiated and differentiated cells. Only ∼5% of polysome-associated lncRNAs pass our stringent thresholds for translation. The translated lncRNA-smORFs exhibit high levels of triplet periodicity and their translational efficiencies are similar to those of protein-coding genes. We can therefore be confident that these are real translation events leading to the production of substantial peptide levels rather than background, spurious events (Bazzini et al., 2014; Guttman et al., 2013; Patraquim et al., 2020; Ruiz-Orera & Alba, 2019). The size distribution of our novel translated ORFs, indicates that the majority are indeed smORFs (<100aa). The general pattern we identified is that dORFs>lncRNA-smORF>uORFs in size. This is consistent with previous studies where a wide range of peptides lengths were discovered (Aspden et al., 2014; Chong et al., 2020). Amino acid composition of these translated smORFs supports the fact they are translated into peptides. However, it does not suggest they are enriched for transmembrane alpha-helices. This is in contrast to the smORFs characterized in *D. melanogaster* (Aspden, 2014).

Overall, we have independent evidence for peptide synthesis for 12/45 lncRNA-smORFs. 8 of these from published mass spectrometry data from Control and RA-differentiated cells SH-SY5Y cells (Brenig et al., 2020). In general, we find smORFs translated in the same treatment as these mass spectrometry datasets detect peptides (7/8). An 18% mass spectrometry detection level may seem low but given the limitations of detecting small peptides by mass spectrometry this represents a substantial level of validation. Two translation events were validated by FLAG tagging transfection assay: LINC01116 and LINC00478 lncRNA-smORFs. The production of 2/45 lncRNA-smORF peptides is corroborated by previous studies in non-neuronal cells. HAND2-AS1 (translated in Control and RA) is translated in human and rodent heart and encodes for an integral membrane component of the endoplasmic reticulum (van Heesch et al., 2019). CRNDE, which is only translated upon differentiation, encodes for a previously characterised nuclear peptide (CRNDEP) (Szafron et al., 2015). The translation of these smORFs in multiple cell types provides substantial support for the production of peptides and their potential function.

We also discovered that 24% of the lncRNA-smORFs we find translated show sequence conservation across the *Hominidae*. This suggests that the other great apes have the potential to translate very similar peptides. This provides additional evidence to indicate that these translation events are not translational noise. Of course, it will be interesting to uncover the function of these small peptides in the future.

Here we have discovered that LINC01116 produces an 87aa peptide that exhibits cytoplasmic localisation, and specifically is detected in near the cell membrane and in neuritic processes. The upregulation of LINC01116 expression upon differentiation as well as the existing evidence of its expression in the developing and adult brain and spinal cord (Consortium, 2013; Lindsay et al., 2016), coupled with the localisation of its peptide, led us to further investigate its potential role in differentiation. Knockdown of LINC01116 upon differentiation appears to impede neurite outgrowth and results in the reduction of the mRNA levels of the noradrenergic marker MOXD1. Our data suggest that LINC01116 is involved in the regulation of neuronal differentiation, consistent with the fact that it is moderately expressed in the developing human forebrain and highly expressed in the developing human midbrain and spinal cord (Lindsay et al., 2016). LINC01116 was previously found to be involved two other cancer models; in the progression of glioblastoma (Brodie et al., 2017) and it is upregulated in gefitinib resistant non-small cell lung cancer cells (Wang He et al., 2020). siRNA knockdown of LINC01116 in both these cell types results in decreased expression of stem-cell markers (Nanog, SOX2 and Oct4) and reduced cell proliferation. This suggests LINC01116 promotes cell proliferation in these systems, indicating that the downstream effects of LINC01116 may vary according to cell type. Interestingly however, knockdown of LINC01116 also inhibited migration of glioma stem cells (GSCs) (Brodie et al., 2017), while overexpression of LINC01116 promoted invasion and migration of gastric cancer cells (Su et al., 2019). This suggests a potential role of LINC01116 in the formation of cell membrane protrusions, which is consistent with the role we have discovered for LINC01116 in neurite development.

Our findings indicate that many lncRNAs are localised in the cytoplasm and play important roles here. Given the large number of lncRNAs associated with polysomes we anticipate that many more lncRNAs will be found to regulate translation. We have identified 121 novel translation events, many of which are regulated during differentiation. Only ∼5% of polysome-associated lncRNAs are translated. The lncRNA-smORFs we discover here represent a general population whose products have not yet been characterized. As we have found for LINC01116, lncRNAs and the small peptides encoded therein have the potential to contribute to important cellular functions and contribute to development and disease.

## Materials and Methods

### Cell culture

Human neuroblastoma SH-SY5Y cells, were cultured in Dulbecco’s Modified Eagle Medium (DMEM 4.5g/L Glucose with L-Glutamine) supplemented with 1% (v/v) Penicillin/Streptomycin and 10% Fetal Bovine Serum (FBS) at 37^°^C, 5% CO_2_. Neural induction commenced at passage 4 and was performed as described previously (Forster et al., 2016; Korecka et al., 2013) with minor alterations. All trans Retinoic Acid (RA, Sigma) was added to cells 24h after plating, at final concentration of 30μM.

### Immunocytochemistry

Cells were seeded on Poly-D-Lysine/mouse laminin coated 12mm round coverslips (Corning BioCoat^™^ Cellware) and fixed with 4% paraformaldehyde (PFA) (Affymetrix) for 20 min at room temperature (RT). A permeabilization step (0.1% Triton-X for 10 min at RT) was performed prior to blocking, followed blocking at RT in blocking buffer (3% BSA, 1XPBS or 5% NGS, 1XPBS and 0.1% Triton-X) for 30 min. Primary antibodies (Antibody Table) were applied in (3% BSA 1X PBS or 0.5% NGS, 1X PBS, 0.1% Triton-X and incubated at RT for 2h or at 4^°^ C overnight. Cells were washed and labelled with Alexa 488, Alexa 555, or Alexa 633 at 1:500 dilution for 2h at RT in 0.5% NGS, 1X PBS, 0.1% Triton-X. Cells were mounted in VECTASHIELD mounting medium, analysed using LSM 700 confocal microscope (Zeiss) ImageJ.

### cDNA synthesis and quantitative Real Time PCR (RT-qPCR)

Equal amounts of RNA (whole cell, nuclear and cytoplasmic lysates) or equal volumes (polysome fractions) were subject to cDNA synthesis, using qScript (Quanta Bio) according to manufacturer’s instructions. qPCR was performed using the CFX Connect^™^ thermal cycler and quantification using SYBR Green fluorescent dye (PowerUp^™^ SYBR Green Master Mix, Thermo Fisher SCIENTIFIC). Primers were designed to anneal to exon-exon junctions, where possible, or to common exons between alternative transcripts (Primer Table). Target mRNA and lncRNA levels were assessed by absolute quantification, by the means of standard curve or relative quantification, using the ΔΔCq method.

### Polysome Profiling

RA was added to SH-SY5Y cells 3 days prior to harvesting. Cells were treated with cycloheximide (Sigma) at 100μg/ml for 3 min at 37^°^C, washed (1X PBS, 100μg/ml cycloheximide) and trypsinised for 5 min at 37^°^C. Subsequently, cells were pelleted, washed (1X PBS, 100μg/ml cycloheximide), and resuspended in ice cold lysis buffer; 50mM Tris-HCl pH8, 150mM NaCl, 10mM MgCl_2_, 1mM DTT, 1% IGEPAL, 100μg/ml cycloheximide, Turbo DNase 24U/μL (Invitrogen), RNasin Plus RNase Inhibitor 90U (Promega), cOmplete Protease Inhibitor (Roche), for 45 min. Cells were then subjected to centrifugation at 17,000 xg for 5 min, to pellet nuclei. Cytoplasmic lysates were loaded onto 18%-60% sucrose gradients (∼70 ⨯10^6^ cells per gradient) at 4^°^C and subjected to ultracentrifugation (121,355 x g_avg_ 3.5h, 4^°^C) in SW-40 rotor. Gradients were fractionated using GRADIENT STATION (Biocomp) and absorbance at 254 nm was monitored using a Biorad detector.

### Poly-Ribo-Seq

∼20% of cytoplasmic lysate was kept for PolyA selection (Total RNA control) and ∼80% was loaded onto 18%-60% sucrose gradients (∼70⨯10^6^ cells per gradient) at 4^°^C and subjected to ultracentrifugation (121,355 ⨯ g_avg_ 3.5h, 4^°^C) in SW-40 rotor. Polysome fractions were pooled from control and from differentiated cells. ∼25% polysomes were kept for polyA selection (Polysome-associated RNA). The remaining 75% was diluted in 100mM Tris-HCl pH8, 30mM NaCl, 10mM MgCl_2_. RNaseI (EN601, 10U/μl 0.7-1U/million cells) was subsequently added incubated overnight at 4^°^C. RNaseI was deactivated using SUPERase inhibitor (200U/gradient) for 5min at 4^°^C. Samples were concentrated using 30 kDa molecular weight cut-off columns (Merck) and loaded on sucrose cushion (1M sucrose, 50mM Tris-HCl pH8, 150mM NaCl, 10mM MgCl_2_, 40U RNase Inhibitor) and subjected to ultracentrifugation at 204,428 ⨯ g_avg_ at 4^°^C for 4h (TLA110). Pellets were resuspended in TRIzol (Ambion, Life Technologies) and processed for RNA purification.

RNA purification from cytoplasmic lysates and RNaseI footprinted samples was performed by Trizol RNA extraction, following manufacturer’s instructions. RNA purification from polysome fractions was performed by isopropanol precipitation, followed by TURBO DNase treatment (Thermofisher) (according to manufacturer’s instructions), acidic phenol/chloroform RNA purification and ethanol precipitation at −80^°^C overnight. RNA concentration was determined by Nano-drop 2000 software. Two rounds of polyA selection from total cytoplasmic lysate and polysome fractions were performed using oligo (dT) Dynabeads (Invitrogen) according to manufacturer’s instructions. PolyA RNA was fragmented by alkaline hydrolysis. 28–34 nt ribosome footprints and 50–80 nt mRNA fragments were gel purified in 10% (w/v) polyacrylamide-TBE-Urea gel at 300V for 3.5h in 1X TBE. Ribosome footprints were subjected to rRNA depletion (Illumina RiboZero rRNA removal kit).

5’ stranded libraries were constructed using NEB Next Multiplex Small RNA Library Prep. Resulting cDNA was PCR amplified and gel purified prior to sequencing. Libraries were subjected to 75bp single end RNA Seq using NextSeq500 Illumina sequencer, High Output Kit v2.5 (75 Cycles) (Next Generation Sequencing Facility, Faculty of Medicine, University of Leeds).

### RNA-Seq data analysis

RNA-Seq reads were trimmed with Cutadapt (Martin, 2011) and filtered with fastq_quality_filter (Hannon, 2010) to filter out the reads of low quality (90% of the read to have a phred score above 20). Filtered reads were mapped (Liao et al., 2013) to human genome reference (the lncRNA GENCODE (Frankish et al., 2019) annotation added to mRNA annotation from UCSC (Haeussler et al., 2019) human genome assembly (hg19) from iGenomes) with Rsubread (Liao et al., 2013) and uniquely mapped reads were reported. Bam file sorting and indexing was performed with SAMtools (Li et al., 2009). Subsequently summarised read counts for all genes were calculated using featureCounts (Liao et al., 2014). For normalization, RPKM values were calculated. Differential expression analysis was conducted between with DESeq2 (Love et al., 2014) based on the two cut-offs p^adj^ < 0.05 and the absolute value of log^2^FoldChange > 1. Gene ontology analysis was performed with GOrilla (***G****ene* ***O****ntology en****RI****chment ana****L****ysis and visua****L****iz****A****tion too*l) (Eden et al., 2009).

### Cytoplasmic/Nuclear fractionation of SH-SY5Y cells

Cells were harvested (as above) and washed with 1X PBS. Cells were lysed in whole cell lysis buffer (see table) (500μL buffer per 10^6^ cells) on ice for 30min. Whole cell lysate aliquots were removed and remainder subjected to centrifugation at 1,600 x g for 8 min to pellet nuclei. Nuclear and cytoplasmic fractions were subjected to two further clearing steps by centrifugation (3,000 x g and 10,000 x g respectively). Nuclei were lysed in RIPA buffer (table). ∼10% of both nuclear and cytoplasmic lysates were used for Western Blot and ∼90% subjected to RNA extraction (ZYMO R1055).

### Western Blot

Samples were diluted in 4X Laemmli sample buffer (Biorad) (277.8 mM Tris-HCI, pH 6.8, 4.4% LDS, 44.4% (v/v) glycerol, 0.02% bromophenol blue), 5% β-mercaptoethanol (Sigma) was added prior to heating at 95^°^C for 5 min and loaded on 10% SDS gels. Gel electrophoresis was performed using the BioRad Mini-PROTEAN 3 gel electrophoresis system (Bio-Rad Laboratories, Hercules, CA, USA). Proteins were transferred to nitrocellulose membranes (Amersham^™^ Protran^™^) and blocked with 5% fat-free milk powder in 1XPBS, 0.05% Tween-20 (Sigma) for 1h at RT. Blots were incubated with primary antibodies overnight (Antibody Table). Blots were then washed in PBS-T and incubated with secondary antibody (anti-mouse HRP) at RT for 2h. Membranes were washed 3 times with PBS-T, prior to application of ECL (Biological Industries). Chemiluminescent signal was detected with Chemi-Doc (BIO-RAD). All membranes were probed for *β*-tubulin as loading control.

### Ribo-Seq analysis

Quality reports of Polysome-associated RNA-Seq and Ribo-Seq data were made using Fastqc (v.0.11.8) (Andrews, 2010). Adaptor sequences were trimmed using Cutadapt (v.1.81) (Martin, 2011) with minimum read length of 25bp, and untrimmed outputs retained for Ribo-Seq reads. Low-quality reads (score <20 for 10% or more of read) were then discarded using FASTQ Quality Filter, FASTX-Toolkit (v.0.0.14) (Gordon, 2010). Human rRNA sequences were retrieved from RiboGalaxy (Michel et al., 2016) and high confidence hg38 tRNA sequences from GtRNAdb Release 17 (Chan & Lowe, 2016). One base was removed from 3’ end of reads to improve alignment quality, reads originating from rRNA and tRNA were aligned and removed using Bowtie2 (v.2.3.4.3) (Langmead & Salzberg, 2012).

The splice aware aligner STAR (v2.6.1b) (Dobin et al., 2012) was used to map remaining reads to human reference genome Release 30 (GRCh38.p12), from Gencode (Frankish et al., 2019). The STAR (v2.6.1b) (Dobin et al., 2012) genome index was built with a sjdbOverhang of 73. Samtools (v.1.9) (Li et al., 2009) was used to create sorted, indexed bam files of the resulting alignments.

Metaplots of aligned Ribo-seq data were generated using create_metaplots.bash script from Ribotaper (v1.3) (Calviello et al., 2016) pipeline. These show the distance between the 5’ ends of Ribo-seq and annotated start and stop codons from CCDS ORFs, allowing the locations of P-sites to be inferred. Read lengths exhibiting the best triplet periodicity were selected for each replicate, along with appropriate offsets (Sup 5).

Actively translated smORFs were then identified using Ribotaper (v1.3) (Calviello et al., 2016). Initially, this requires an exon to contain more than 5 P-sites in order to pass to quality control steps. Identified ORFs were then required to have a 3-nt periodic pattern of Ribo-seq reads, with 50% or more of the P-sites in-frame. In the case of multiple start codons, the most upstream in-frame start codon with a minimum of five P-sites in between it and the next ATG was selected.

ORFs for which >30% of the Ribo-seq coverage was only supported by multimapping reads were also subsequently filtered. For a smORF to be considered actively translated in a condition, we required that it be identified in at least two of the three biological replicates for the condition.

### Translation efficiency calculation

Translational efficiency (TE) was estimated for all translated ORFs in each condition, where TE was equal to the mean number of P sites per ORF, normalised by the median P sites per ORF per replicate, divided by the mean number of RNA sites per ORF, normalised by the median RNA sites per ORF per replicate.

### Differential translation efficiency analysis

Differential translation efficiency analysis was performed using the ΔTE (Chothani et al., 2019) alternate protocol. However, our analysis was based on read counts from all identified translated ORFs from the Ribo-seq and polysome-associated RNA-seq, not the whole CDS as described in the protocol.

### Amino acid composition

For each of our ORF sets (protein coding, lncRNA-smORF, uORF and dORFs), the average amino acid compositions were calculated. Random control expected frequencies were taken from King and Jukes (King & Jukes, 1969).

### Mass spectrometry analysis

Two published SH-SY5Y cell mass proteomics datasets were analysed: PXD010776 (Murillo et al., 2018) and PXD014381 (Brenig et al., 2020). Binary raw files (*.raw) were downloaded from PRIDE then converted to human-readable MGF format using ThemoRawFileParser (Hulstaert et al., 2020). The amino acid sequences of our translated uORFs, dORFs and lncRNA-smORFs were added to the whole *Homo sapiens* proteome dataset (20,379 entries) downloaded from UniProtKB (Bateman et al., 2019) on Nov/2019. The new FASTA file was then used as a customized database on Comet (v2019.01.2) (Eng et al., 2013) search engine runs that scanned all MS/MS files (*.mgf) against it.

Default settings in Comet were used with the following exceptions according to the MS/MS data type. iTRAQ-4plex (PXD010776): decoy_search = 1, peptide_mass_tolerance= 10.00, fragment_bin_tol = 0.1, fragment_bin_offset = 0.0, theoretical_fragment_ions = 0, spectrum_batch_size = 15000, clear_mz_range = 113.5-117.5, add_Nterm_peptide = 144.10253, add_K_lysine = 144.10253, minimum_peaks = 8. Label-free (PXD014381): decoy_search = 1, peptide_mass_tolerance = 10.00, fragment_bin_tol = 0.02, fragment_bin_offset = 0.0, theoretical_fragment_ions = 0, spectrum_batch_size = 15000.

CometUI (Eng et al., 2013) was employed for assessing Comet’s *.xml output files and setting a false discovery rate (FDR) threshold of 10% per peptide identification. This FDR threshold was selected due to expected low abundance levels of the target smORFs.

### Conservation analysis

Protein, cDNA and ncRNA sequence data for *H*.*sapiens, P*.*abelii, P*.*paniscus, P*.*troglodytes, G*.*gorilla, N*.*leucogenys*, and *M*.*musculus* were obtained from Ensembl (release 100, (Yates et al., 2020).

A sequence homology search was performed using the 45 translated lncRNA peptide sequences, against each non-human species protein database using BLASTp (e value - 0.001) (Altschul et al., 1990). As these are small peptides, results were filtered to remove anything with <75% identity, unless a result(s) was the lowest e value hit for a given query in each species. Results were returned for 12 lncRNA peptides, and manually cross validated using Ensembl Genome Browser and multiple sequence alignments performed using ClustalOmega (Sievers et al., 2011), run using default parameters using the msa package in R (Bodenhofer et al., 2015), to give 8 peptides with evidence of sequence conservation.

The transcript sequences of the 45 translated lncRNAs were searched against transcriptome databases created by combining the cDNA and ncRNA data for each species, using BLASTn (e value - 0.001) (Altschul et al., 1990). Results of this BLAST were used to filter the initial BLAST databases. ORF portions of the 45 translated lncRNAs were extracted, and searched against these filtered databases using BLASTn (e value - 0.001)(Altschul et al., 1990).

It was confirmed that all the lncRNA ORFs returned their genes of origin in *H*.*sapiens*. The remaining species returned results for 18 of the lncRNA ORF queries. These were cross validated as above, resulting in 14 lncRNA ORFs with evidence of sequence conservation based on transcript sequences.

A final search was performed using the 45 translated lncRNA peptide sequences against the transcriptome databases using tBLASTn (e value - 0.001) (Altschul et al., 1990). To reduce redundancy, results were filtered to select the transcripts(s) with lowest e value for each gene. It was confirmed that all ORFs returned their genes of origin in *H*.*sapiens*.

The remaining species returned results for 21 of the lncRNA peptide queries. As some queries had many spurious results, they were further filtered to select the transcripts(s) with lowest e value for each query in each species. These were cross validated as above, resulting in 16 lncRNA peptides with evidence of sequence conservation based on transcript sequences.

We combined evidence from all these approaches into a final dataset consisting of 17 lncRNA small ORFs with evidence of conservation.

### Plasmid construction/Cloning

5’-UTRs and CDSs of putative smORFs (lacking stop codon) were generated by PCR (Table 3), using NEB High Fidelity DNA Polymerase (Q5). C-terminal 3xFLAG tag was incorporated within the reverse primer (Table 3 marked with green) by PCR and products were cloned into NheI and EcoRV restriction sites (Table 3: marked with blue) of pcDNA3.1-hygro vector (Addgene, kindly provided by Mark Richards-Bayliss group, University of Leeds). Start codon mutations were generated by site directed mutagenesis (Q5 Site directed mutagenesis kit, NEB).

**Table 3:**
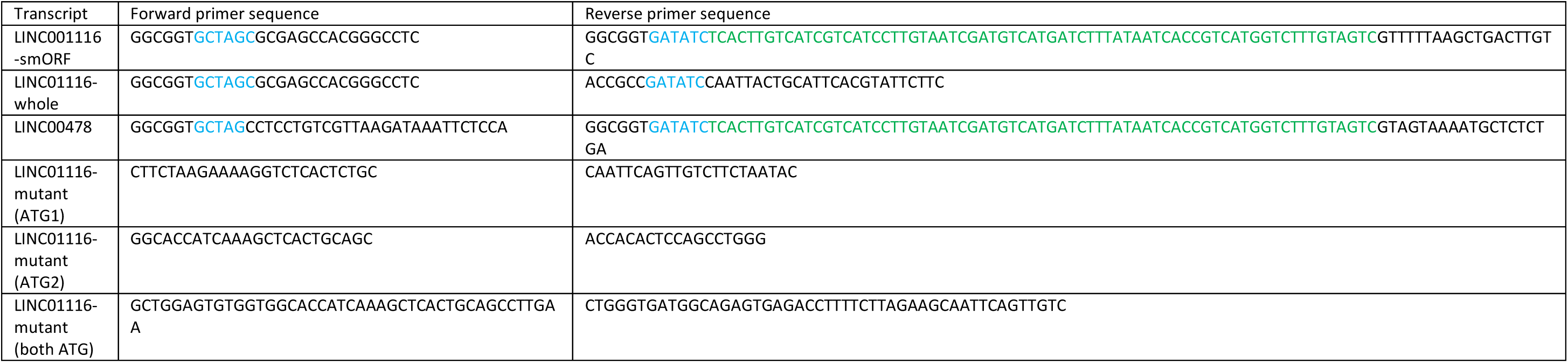
Cloning Primers.

**Table 4:** Antibodies.

| Protein | Species | Company | Usage (ICC/WB) | Dilution |
| --- | --- | --- | --- | --- |
| betaIII-tubulin (TuJ1) | rabbit | ProteinTech | ICC | 1:50 |
| ki67 | mouse | Dako-Agilent | ICC | 1:100 |
| c-Fos | rabbit | Santa Cruz | ICC | 1:50 |
| Tubulin B | mouse | DSHB | both | 1:5000 for WB,<br>1:200 for ICC |
| NXF1 | mouse | abcam | WB | 1:5000 |
| H3K27me3 | mouse | abcam | WB | 1:1000 |
| FLAG | mouse | Sigma | both | 1:1000 |
| hnRNPK | rabbit | abcam | ICC | 1:100 |
| anti-mouse IgG-HRP | goat | New England Biolabs | WB | 1:5000 |
| anti-rabbit IgG-HRP | goat | New England Biolabs | WB | 1:5000 |
| Alexa anti-mouse IgG-488 | goat | Thermofisher | ICC | 1:500 |
| Alexa anti-rabbit IgG-488 | goat | Thermofisher | ICC | 1:500 |
| Alexa anti-rabbit 633 | goat | Thermofisher | ICC | 1:500 |

### Transfections and Microscopy

Plasmid transfections were performed using Lipofectamine 3000 (Thermo) following the manufacturer’s instructions. After 48 h, the cells were fixed for 20 min with 4% paraformaldehyde, washed with 1X PBS, 0.1% Triton X–100 (PBS-T) and processed for immunocytochemistry as previously described. Imaging was conducted using EVOS fluorescent microscope.

### siRNA knockdown

siRNA knockdown was performed using Lincode siRNA SMARTpool (Dharmacon) (LINC01116 transcript: R-027999-00-0005 SMARTpool). Lincode Non-targeting Pool (D-001810-10) was used as scrambled control. Cells were seeded in 24-well plates (10^5^ cells/well,) and siRNA where transfected using RNAiMAX lipofectamine (ThermoFisher) as per manufacturer’s instructions.

### General statistics and plots

Statistical analyses were performed in R (R Core Team, 2019), using packages including stringr (Wickham, 2019), dplyr (Wickham, 2017), tidyr (Wickham, 2017), protr (Xiao et al., 2015) ggplot (Wickham, 2016), ggstatsplot (Patil, 2018), knitr (Xie, 2020), seqinr (Charif D, 2007), ggbeeswarm (Clarke & Sherrill-Mix, 2017), and EnhancedVolcano (Blighe et al., 2018)

Experimental values from independent samples with equal variances, were assessed using 2-tailed unpaired Student’s t-test. The results are shown as mean±SEM values of 3 independent replicates. The exact P values are described and specified in each figure legend. P values < 0.05 were considered statistically significant.

## Data availability

Poly-Ribo-Seq datasets will be deposited in GEO.

## Acknowledgements

We wish to thank Dr Eric Hewitt for providing the SH-SY5Y cells; Dr Iosifina Sampson and Dr Mark Richards (Bayliss group) for providing HEK293 cells and mammalian vectors. We thank the Next Generation Sequencing facility, at St James University Hospital, Leeds, UK for performing Next Generation Sequencing and Imaging Facility, Faculty of Biological Sciences, University of Leeds, UK for their assistance in confocal microscopy. Parts of this work were undertaken on ARC3, part of the High Performance Computing facilities at the University of Leeds, UK.

## Author contributions

KD designed and performed experiments, acquired, analysed and interpreted data, drafted and revised the manuscript. IB designed and performed experiments, acquired, analysed and interpreted data, drafted and revised the manuscript. DW analysed, interpreted data and revised the manuscript. SC acquired and analysed data, and revised the manuscript. AB acquired and analysed data, and revised the manuscript. EJRV analysed data and revised the manuscript. MOC helped interpret portions of the data and critique the output for important intellectual content, and revised the manuscript. JD helped design experiments, interpret portions of the data and critique the output for important intellectual content, and revised the manuscript. AW helped design experiments, interpret portions of the data and critique the output for important intellectual content, and revised the manuscript. JLA conceived the work, interpreted data, drafted and revised the manuscript.

---

**Sup Figure 1:**
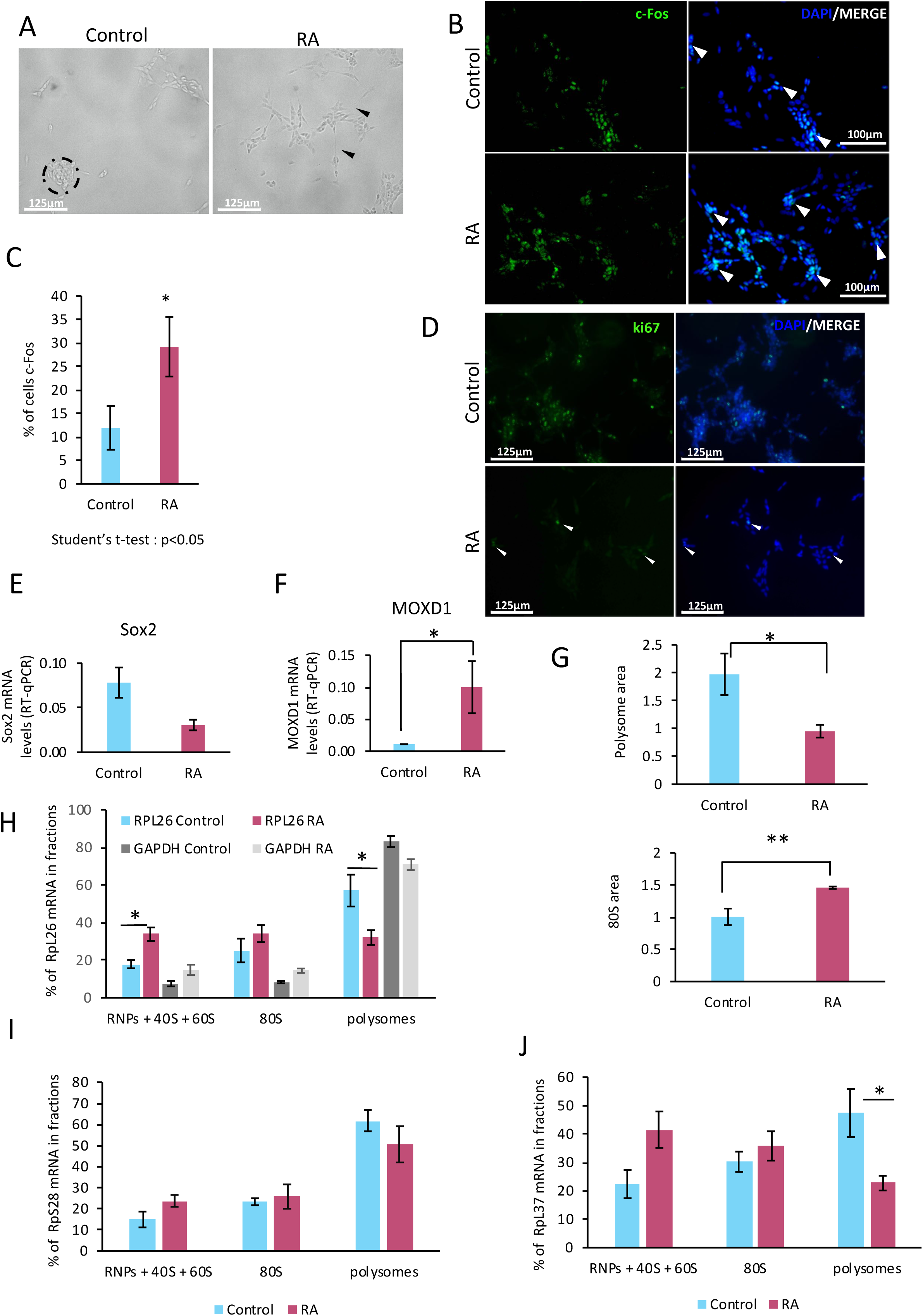
Differentiation of SH-SY5Y results in a reduction in the level of active translation. (A) Light microscopy reveals that undifferentiated SH-SY5Y cells grow in clumps and upon neural induction with RA they extend their neurites. (B) lmmunofluorescence indicates c-Fos differentiation marker is expressed at higher levels in differentiated cells. (C) Quantification of c-Fos+ cells in Control and RA (Student’s t-test, n=1000, <0.05). (D) lmmunofluorescence shows ki67 proliferation marker expression is reduced upon differentiation. (E) RT-qPCR of pluripotency/proliferation marker SOX2 shows reduced levels upon differentiation. (F) RT-qPCR of the differentiation marker MOXD1 indicates differentiation towards outer radial glia neural precursors (Student’s t-test, n=3, p<0.05). (G) Quantification of polysome (Student’s t-test, n=3, p<0.05) and monosome (Student’s t-test, n=3, p<0.01) areas upon differentiation. (H) RT-qPCR of RPL26 mRNA distribution (Control; orange, RA; green) (with GAPDH mRNA controls; Control; light grey, RA; dark grey), in RNPs and ribosomal subunits (RNPs,40S, 60S), monosome (80S) and polysomes (Student’s t-test, n=3, p<0.05). RT-qPCR across gradients of RPS28 (I) and RPL37 (J) mRNAs respectively (Student’s t-test, n=3, p<0.05).

**Sup Figure 2:**
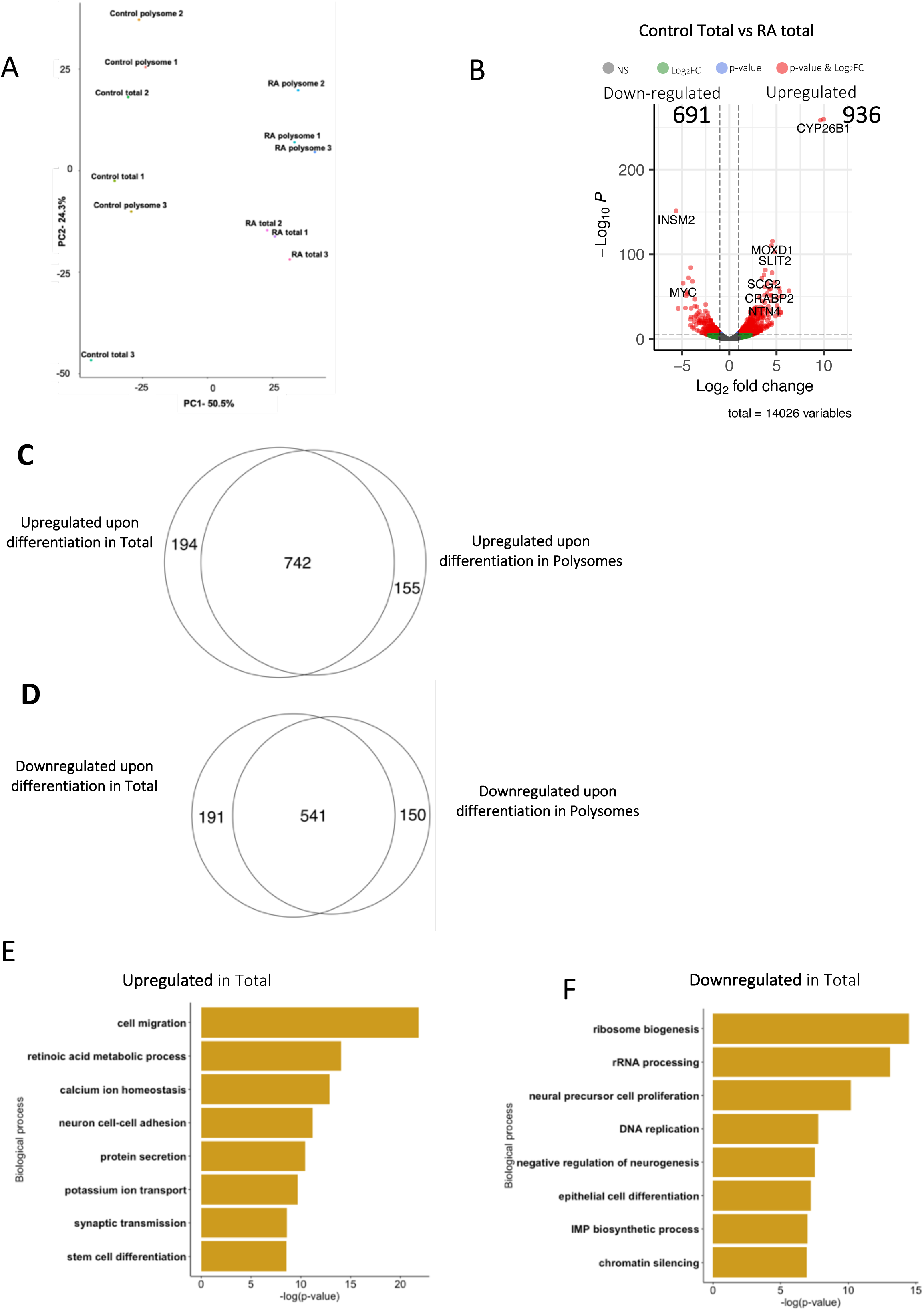
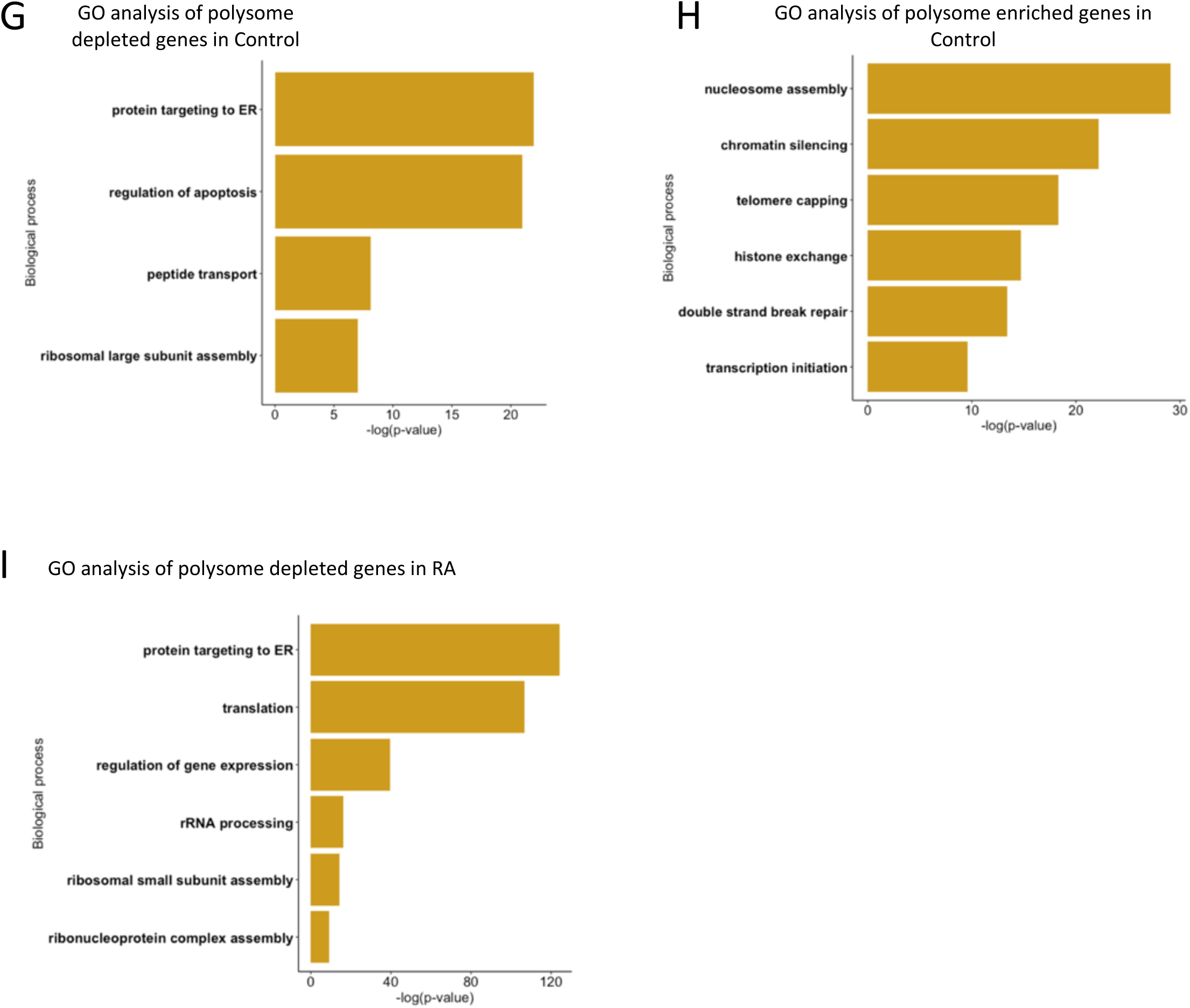
Differentiation results in global RNA changes. (A) PCA analysis of the RNA-Seq datasets shows that Control samples are more similar to each other than to RA treated samples. Control samples cluster separately from RA samples, indicating significant variation in gene expression between Control and differentiated cells. RA total samples cluster separately from RA polysome samples, whereas Control samples do not show different clustering, indicating that the variation in gene expression between RA total and RA polysome is greater than the variation between Control total and Control pololysome. (B) Volcano plot depicting the differentially expressed protein-coding genes between Total Control and Total RA populations; 936 protein-coding mRNAs are upregulated and 691 downregulated upon differentiation (log_2_ fold-change cut-off=1, p^adj^<0.05). Venn diagrams of overlap between those mRNAs identified as up (C) or down (D) regulated between Total and Polysome populations. GO term analysis for total cytoplasmic protein-coding genes (E) upregulated in and (F) downregulated. GO term analysis for protein-coding genes; (G) depleted from polysomes in Control, (H) enriched in polysomes in Control and (I) depleted from polysomes in RA.

**Sup Figure 3:**
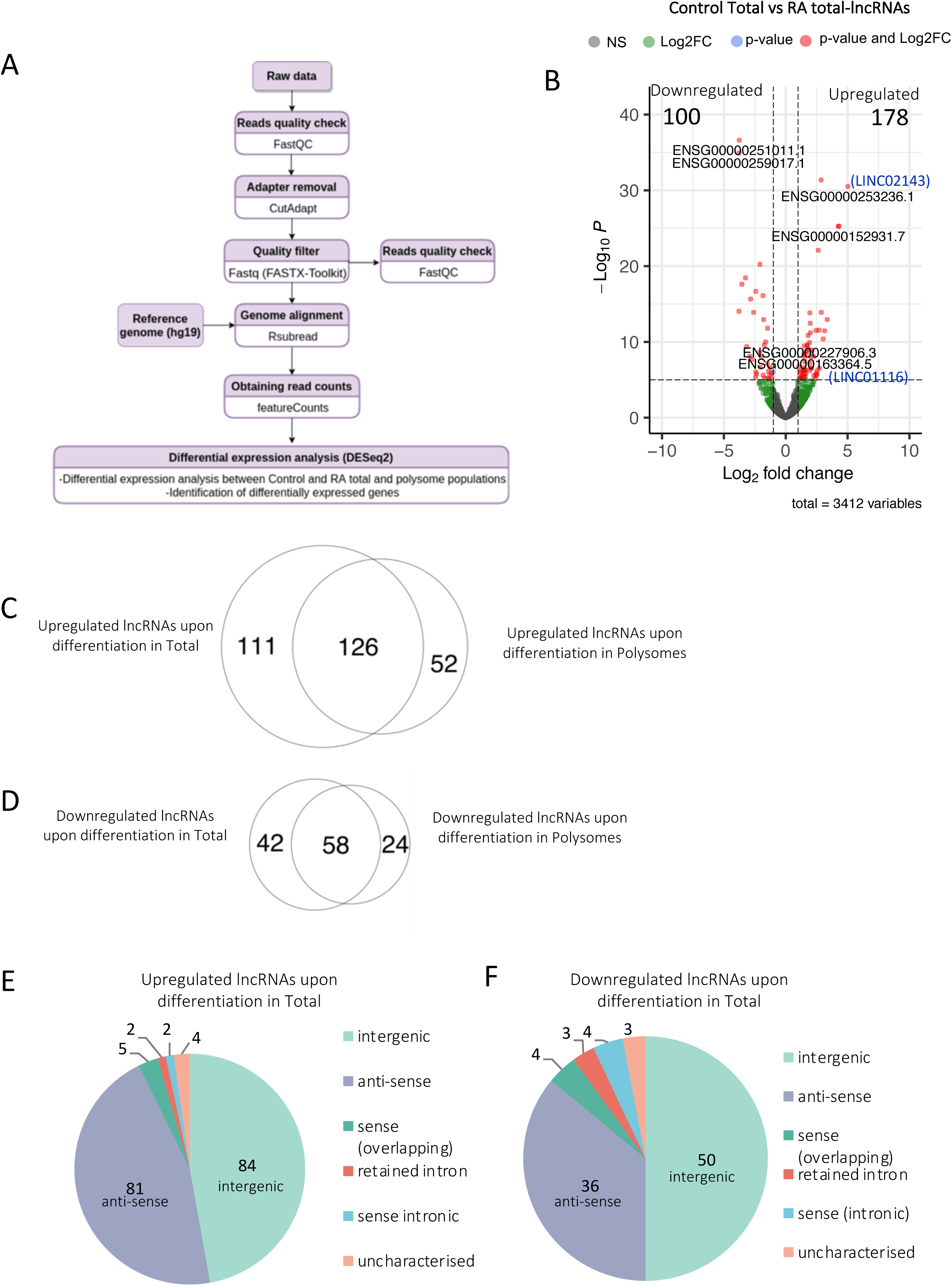
Regulation of lncRNA expression and polysome association. (A) Pipeline of lncRNA differential expression analysis. (B) Volcano plot displaying the significantly differentially expressed lncRNAs (labelled by their geneIDs) between Control and RA populations (Total) with log_2_ fold-change cutoff=1, p^adj^<0.05. Venn diagrams of overlap in differentially expressed lncRNAs between Total and Polysomes datasets for (C) upregulated lncRNAs and (D) downregulated lncRNAs upon differentiation. Pie charts showing breakdown by lncRNA type for lncRNAs (E) upregulated and (F) downregulated upon differentiation (Polysome) (dark purple: intergenic; magenta: anti-sense; purple: sense -overlapping; dark pink: retained intron; light pink: sense-intronic; white: uncharacterized; bright pink: NMD target).

**Sup Figure 4:**
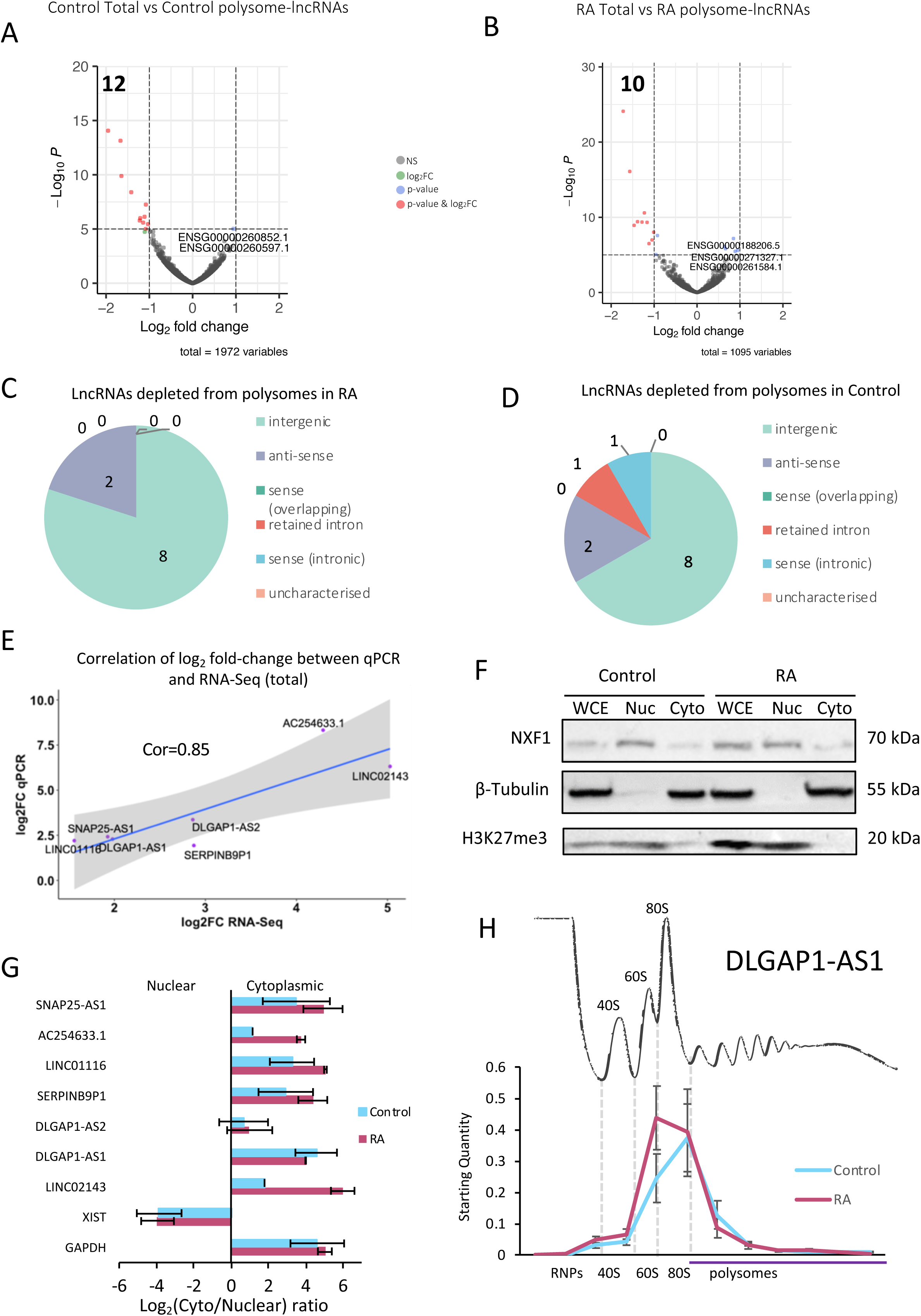
Regulation of lncRNA expression and polysome association. Volcano plots displaying the significantly differentially localised lncRNAs (labelled by their geneIDs) between Total and Polysome populations in (A) Control and (B) RA with log_2_ fold-change cutoff=1, p^adj^<0.05. Pie charts of lncRNA types present in different populations (C) depleted from polysomes in RA and (D) depleted from polysomes in Control (intergenic; anti-sense; sense-overlapping; retained intron; sense-tronic; uncharacterized; NMD target). (E) Correlation of RT-qPCR data for fold-change for lncRNAs with RNA-Seq analysis (LINC01116, LINC02143, SNAP25-AS1, DLGAP1-AS1, DLGAP1-AS2, SERPINB9P1 and AC254633.1). (F) Western blot confirming cellular fractionation with nuclear marker (H3K27me3) and cytoplasmic markers (beta-tubulin). (G) LncRNAs of interest that are induced are specifically localised to cytoplasm as shown by subcellular localisation RT-qPCR for Control and RA samples. XIST lncRNA was used as a nuclear and GAPDH mRNA as a cytoplasmic positive control (n=3, SE is plotted). (H) RT-qPCR of lncRNAs across sucrose gradient fractions indicates that DLGAP1-AS2 is found in 80S and small polysome fractions during differentiation and fractions both in control and RA treated cells (n=3, SE is plotted) On average, 63% of the transcripts is detected in the polysome fractions in Control and 49% upon differentiation.

**Table 1:** Translation of small ORFs. Number of ORFs detected as translated in Poly-Ribo-Seq

| Translated ORFs | Control | RA | Overlap | Total |
| --- | --- | --- | --- | --- |
| Protein-Coding ORFs | 16,282 | 12,745 | 10,014 | 19,013 |
| uORFs | 56 | 27 | 12 | 71 |
| dORFs | 4 | 1 | 0 | 5 |
| lncRNA-smORFs | 28 | 23 | 6 | 45 |

**Sup Figure 5:**
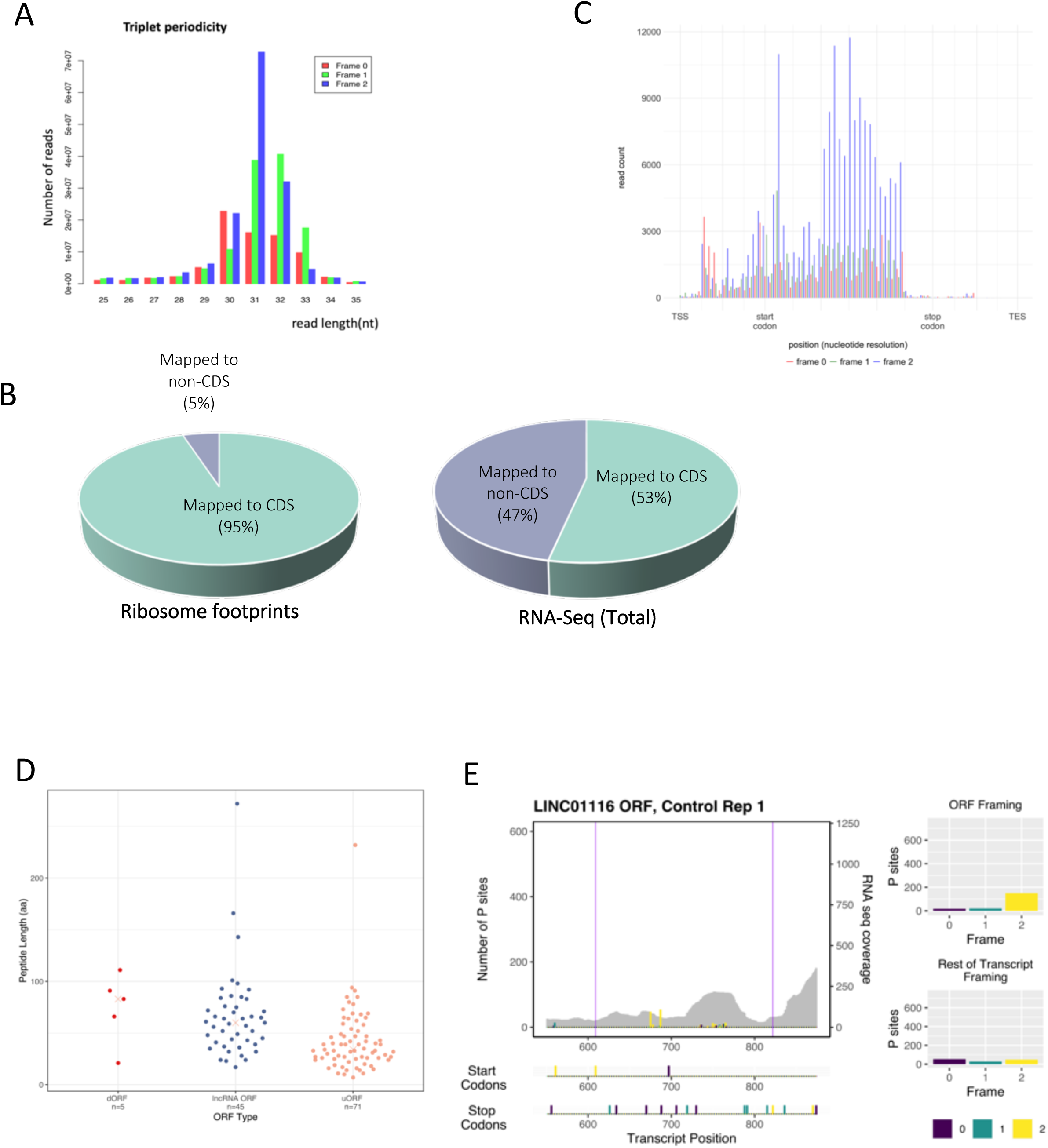
Translation of lncRNA smORFs. (A) Triplet periodicity plot for 25-35 nt reads from RA sample, replicate 3. (B) Pie charts of numbers of reads mapping to CDSs and elsewhere for ribosome footprints and RNA-Seq (Total). Values are mean of three Control replicates. (C) Metagene analysis for 33 nt reads from 33 nt reads from replicate x, Control sample. (D) Length distribution of translated ORFs inlnc RNAs, dORFs and uORFs (in codons), results from Control and RA combined. (E) Example Poly-Ribo-Seq profile for LINC01116 in control treatment. RNA-Seq (Polysome) reads are grey and ribosome P sites in red, green and blue according to frame. Purple lines mark beginning and end of translated smORF. All possible start and stop codons are indicated below. Framing within and outside translated smORF shown on left.

**Sup Figure Table 1.**
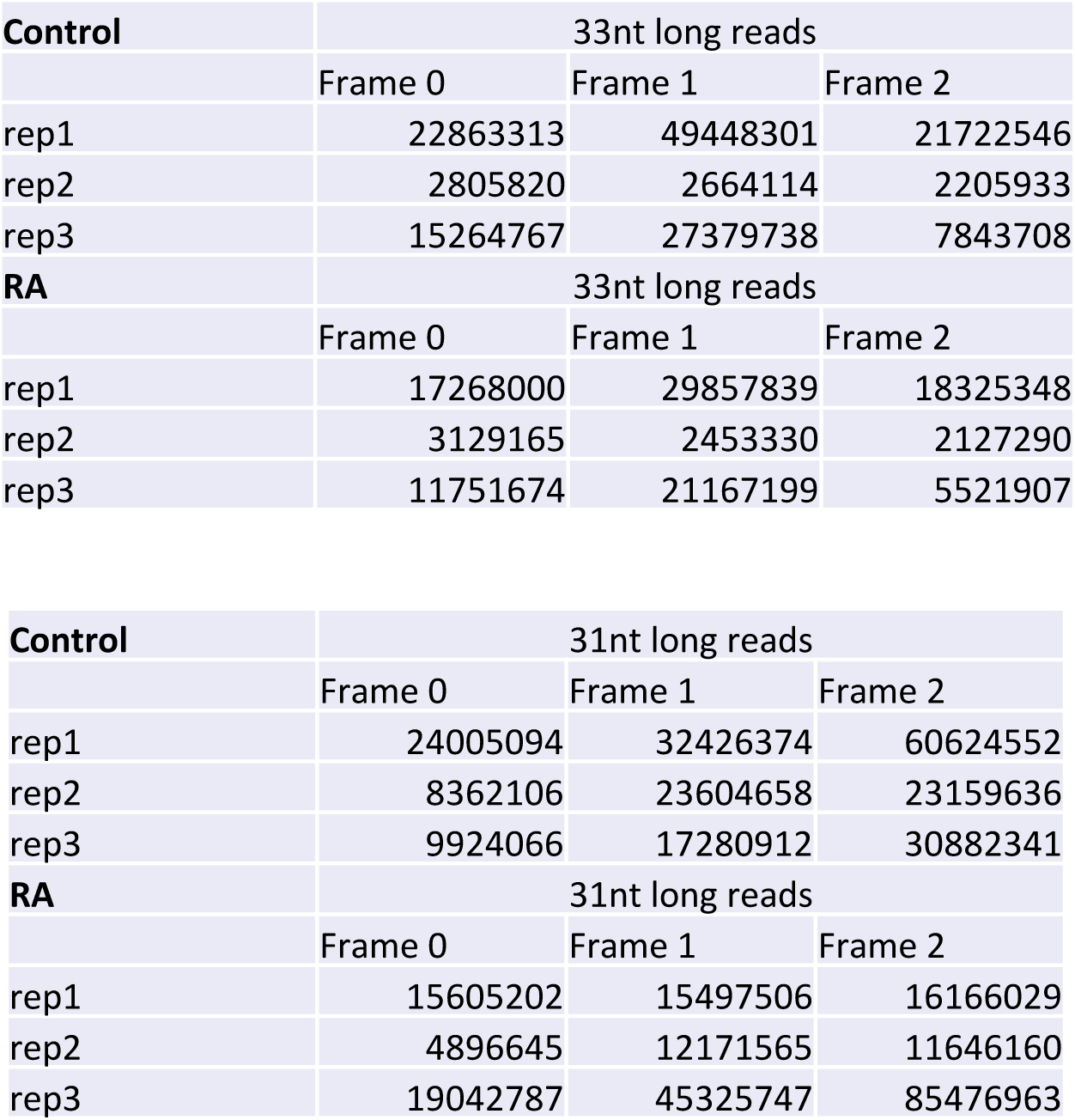
Summary of triplet periodicity data for all 3 replicates, Control and RA, for 31 nt and 33 nt footprint lengths.

**Sup Figure 6:**
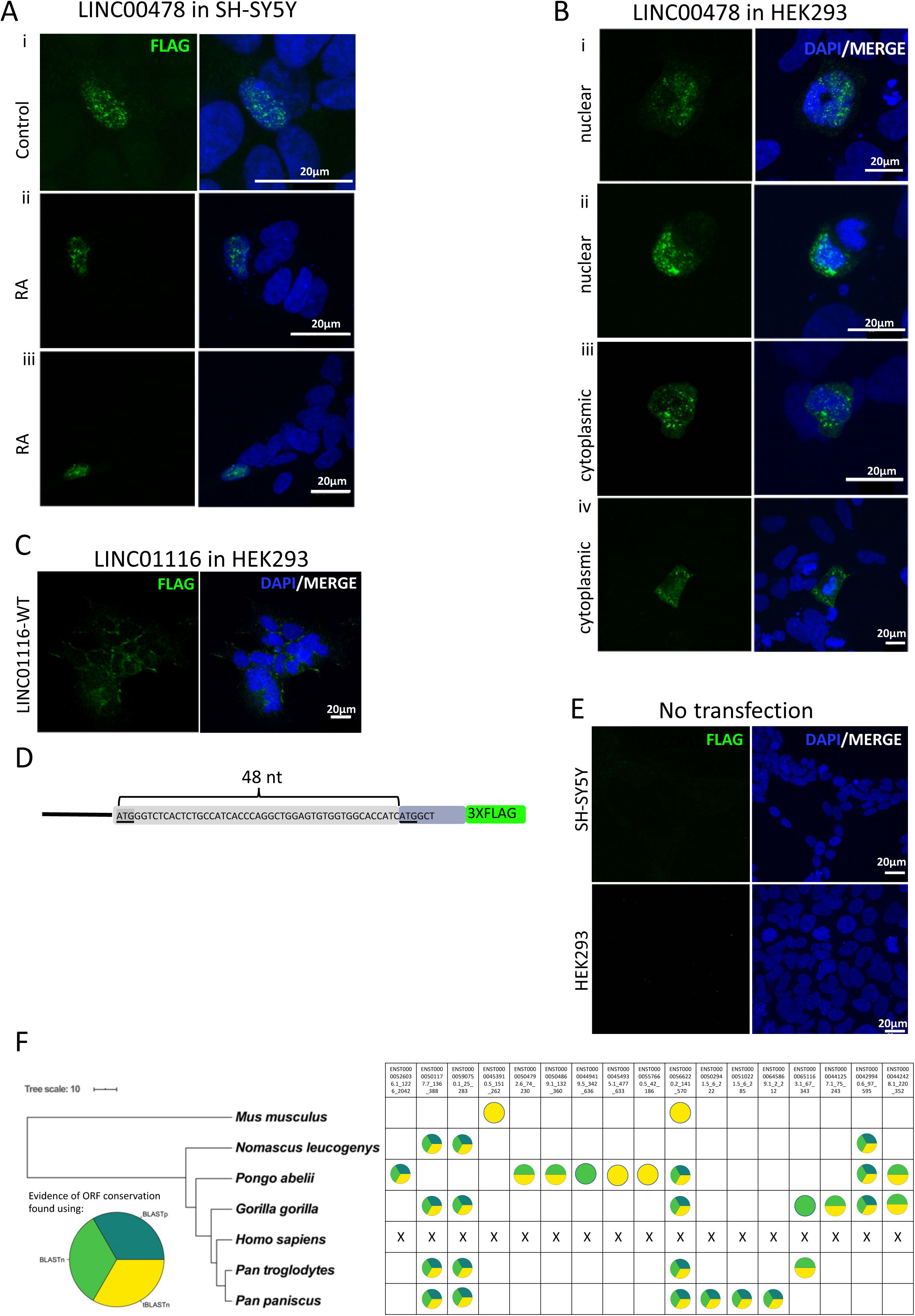
Peptide production from smORFs in lncRNAs. (A) Representative confocal images of FLAG-tagged LINC00478 peptide in (i) Control and (ii-iii) RA treated SH-SY5Y cells. LINC00478 exhibits nuclear distribution. (B) Representative confocal images of transfections of FLAG-tagged LINC00478 smORF in HEK293 cells; showing (i-ii) nuclear and (iii) cytoplasmic distribution, greenisFLAG, and blueis DAPI (scale bar is 20μm). (C) Representative confocal images of FLAG-tagged LINC01116 smORF (i) WT showing cytoplasmic distribution near cell membrane and filopodia. Green is FLAG and blue is DAPI (scale bar is 16μm). (D) schematic of LINC01116 smORF with2 start codons annotated.(E) Negative control (no transfection) in (i) SH-SY5Y and (ii) HEK293 cells; green is FLAG and blue is DAPI (scale bar is 20μm). (F) Phylogram with lncRNA-smORFs for which evidence of sequence conservation were found represented as circles, coloured according to how sequence conservation was identified. Each lncRNA-smORF with evidence of sequence conservation is shown along with which species shown conservation. Phylogram built in iTOL (Letunic, I. et al .2006) using data from TimeTree (Kumar, S.,et al. 2017),scale in 10 MYA along.

**Sup Figure 7:**
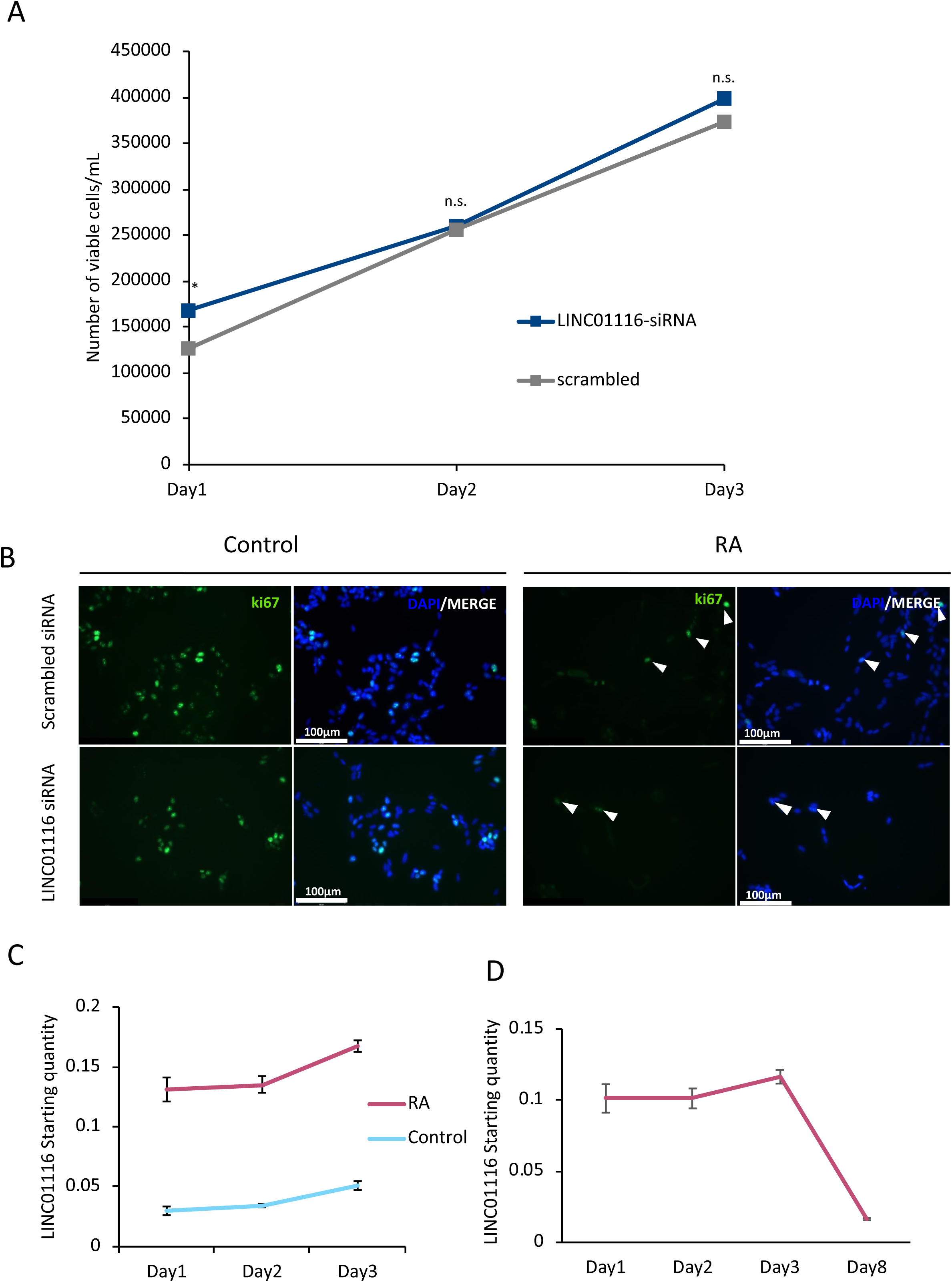
LINC01116 expression is upregulated early on upon differentiation and does not affect cell cycle progression. (A) LINC01116 knockdown is not cytotoxic, as shown by cell viability assay (N=3 biological replicates, n=2 technical duplicates per replicate, student’s t-test p>0.05). (B) Representative immunofluorescence of Control and RASH-SY5Y cells, transfected with LINC01116 or scrambled siRNA, after staining for proliferation marker ki67 (arrowheads mark ki67+ cells) at day 3 post differentiation (N=3 biological replicates, n>100 measurements, student’s t-testp>0.05). Scale bar = 100μm(C) Expression of LINC01116 lncRNA is upregulated from first day post-differentiation as shown by RT-qPCR (N=3 biological replicates, student’s t-testp<0.05).(D) LINC01116 expression levels, in phase 1 and phase 2 differentiated SH-SY5Y cells, measured by RT-qPCR. LINC01116 levels decrease ∼6-fold during phase 2 differentiation (n=3, standard deviation is plotted).

